# An expedited approach towards the rationale design of non-covalent SARS-CoV-2 main protease inhibitors with in vitro antiviral activity

**DOI:** 10.1101/2020.12.19.423537

**Authors:** Naoya Kitamura, Michael Dominic Sacco, Chunlong Ma, Yanmei Hu, Julia Alma Townsend, Xiangzhi Meng, Fushun Zhang, Xiujun Zhang, Adis Kukuljac, Michael Thomas Marty, David Schultz, Sara Cherry, Yan Xiang, Yu Chen, Jun Wang

## Abstract

The main protease (M^pro^) of SARS-CoV-2 is a validated antiviral drug target. Several M^pro^ inhibitors have been reported with potent enzymatic inhibition and cellular antiviral activity, including GC376, boceprevir, calpain inhibitors II and XII, each containing a reactive warhead that covalently modifies the catalytic Cys145. In this study, we report an expedited drug discovery approach by coupling structure-based design and Ugi four-component (Ugi-4CR) reaction methodology to the design of non-covalent M^pro^ inhibitors. The most potent compound **23R** had cellular antiviral activity similar to covalent inhibitors such as GC376. Our designs were guided by overlaying the structure of SARS-CoV M^pro^ + ML188 (R), a non-covalent inhibitor derived from Ug-4CR, with the X-ray crystal structures of SARS-CoV-2 M^pro^ + calpain inhibitor XII/GC376/UAWJ247. Binding site analysis suggests a strategy of extending the P2 and P3 substitutions in ML188 (R) to achieve optimal shape complementary with SARS-CoV-2 M^pro^. Lead optimization led to the discovery of **23R**, which inhibits SARS-CoV-2 M^pro^ and SARS-CoV-2 viral replication with an IC_50_ of 0.31 μM and EC_50_ of 1.27 μM, respectively. The binding and specificity of **23R** to SARS-CoV-2 M^pro^ were confirmed in a thermal shift assay and native mass spectrometry assay. The co-crystal structure of SARS-CoV-2 M^pro^ with **23R** revealed the P2 biphenyl fits snuggly into the S2 pocket and the benzyl group in the α-methylbenzyl faces towards the core of the enzyme, occupying a previously unexplored binding site located in between the S2 and S4 pockets. Overall, this study revealed the most potent non-covalent SARS-CoV-2 M^pro^ inhibitors reported to date and a novel binding pocket that can be explored for M^pro^ inhibitor design.

---

The COVID-19 pandemic had a significant impact on global economy and public health, and there is an urgent need for therapeutic interventions. Encouraging progress has been made in developing mRNA vaccines including the Pfizer BNT162b2 and Moderna mRNA-1273. For small molecule antivirals, the viral polymerase inhibitor remdesivir gained FDA approval on Oct 22^nd^ 2020. The combination therapy of remdesivir with a Janus kinase (JAK) inhibitor baricitinib also received the FDA emergency use authorization^1^. Among the other drug targets being explored at different stages of preclinical and clinical developments^2^, the viral main protease (M^pro^), also called 3-chymotrypsin-like protease (3CL^pro^), is one of the most extensively explored high profile antiviral drug targets^3^. M^pro^ is a cysteine protease encoded in the viral polyprotein as non-structural protein 5 (Nsp5) that cleaves the viral polyproteins pp1a and pp1ab at more than 11 sites. Despite its multiple proteolytic sites, M^pro^ was shown to have a high substrate specificity of glutamine at the P1 position^4^. As such, the majority of the reported M^pro^ inhibitors were designed to contain a 2-pyrrolidone at the P1 substitution as a mimetic of the glutamine in the substrate^5^. Most advanced M^pro^ inhibitors including PF-07304814^6^, GC376^7,8^ and 6j^9^ all belong to this category. PF-07304814, an α-hydroxyl ketone prodrug, is being developed by Pfizer, which has optimal pharmacokinetic properties in humans and recently entered human clinical trials^6^. GC376 has *in vivo* antiviral efficacy in treating cats infected with lethal feline infectious peritonitis virus^10,11^. Recently, the GC376 analog 6j was shown to protect mice from MERS-CoV infection^9^. These promising results highlight the translational potential of M^pro^ inhibitors as potent SARS-CoV-2 antivirals and validate M^pro^ as an antiviral drug target for coronaviruses.

Drug discovery is a lengthy process, which involves iterative cycles of design, synthesis, and pharmacological characterization. In the event of COVID-19 pandemic, an expedited approach with a fast turnover of this development cycle is highly desired. Using SARS-CoV-2 M^pro^ as a drug target, we report herein a fast-track drug discovery approach by coupling structure-based drug design and the Ugi four-component reaction (Ugi-4CR) methodology to the design of non-covalent M^pro^ inhibitors. Specifically, through a screening of a focused library of protease inhibitors, we recently discovered several non-canonical SARS-CoV-2 M^pro^ inhibitors including boceprevir, and calpain inhibitors II and XII^7^. These inhibitors differ from classic M^pro^ inhibitors such as GC376 in that their P1 substitution does not contain a glutamine mimetic. The co-crystal structures of calpain inhibitors II and XII with SARS-CoV-2 M^pro^ revealed a critical hydrogen bond between the methionine side chain from calpain inhibitor II and pyridinyl substitution from calpain inhibitor XII with the H163 side chain imidazole located at the S1 pocket^8^. Similarly, the carbonyl from the pyrrolidone in GC376 also forms a hydrogen bond with the H163 side chain imidazole^7^. Given the importance of this hydrogen bond with H163 for the high affinity binding of inhibitors to SARS-CoV-2 M^pro^, we hypothesize that non-covalent inhibitors without a reactive warhead targeting the Cys145, but retain the hydrogen bond capacity with H163 can be designed as potent SARS-CoV-2 M^pro^ inhibitors. In this study, we report the structure-based design of non-covalent M^pro^ inhibitors based on the overlaying structures of SARS-CoV or SARS-CoV-2 M^pro^ in complex with existing inhibitors or the peptide substrate. The design was based on the scaffold of ML188 (R)^12^, a non-covalent SARS-CoV M^pro^ inhibitor, which similarly contains a pyridinyl in the P1 substitution as calpain inhibitor XII. The overlaying structures revealed a strategy of extending the P2 and P3 substitutions in ML188 (R) to occupy the extra space in the S2 and S3/S4 pockets of SARS-CoV-2 M^pro^ in order to increase the binding affinity. The most potent inhibitor from this study **23R** showed enzymatic inhibition and cellular antiviral activity similar to the covalent inhibitor GC376. Its mechanism of action was characterized in the thermal shift-binding assay, native mass spectrometry binding assay, and enzyme kinetic studies. An X-ray crystal structure of SARS-CoV-2 M^pro^ in complex with **23R** was solved, revealing a previously unexplored binding site in between the S2 and S4 pockets. Overall, this study led to the identification of the most potent non-covalent M^pro^ inhibitor **23R** with potent enzymatic inhibition and *in vitro* cellular antiviral activity with a novel mechanism of action.

## RESULTS

### Rational design of non-covalent SARS-CoV-2 M^pro^ inhibitors

Among the non-canonical SARS-CoV-2 M^pro^ inhibitors we recently discovered, calpain inhibitor XII has an unexpected binding mode showing an inverted conformation in the active site^8^ (Fig. 1a). Instead of projecting the norvaline and leucine side chains into the S1 and S2 pockets as one would expect from its chemical structure, the pyridinyl substitution snuggly fits in the S1 pocket and forms a hydrogen bond with the H163 imidazole (Fig. 1a). This hydrogen bond is essential, as replacing the pyridine with benzene led to an analog UAWJ257 with a significant loss of enzymatic inhibition^8^. Examining the X-ray crystal structures of SARS-CoV and SARS-CoV-2 M^pro^ in the PDB database revealed another compound ML188 (R)^12^, which shares a similar binding mode with calpain inhibitor XII. ML188 (R) is a non-covalent SARS-CoV M^pro^ inhibitor derived from a high-throughput screening hit^12^. The pyridinyl from ML188 (R) similarly fits in the S1 pocket and forms a hydrogen bond with the H163 side chain imidazole (Fig. 1b). In addition, the furyl oxygen and its amide oxygen both form a hydrogen bond with the G143 main chain amide amine. ML188 (R) was reported to inhibit the SARS-CoV M^pro^ with an IC_50_ value of 1.5 ± 0.3 μM and the SARS-CoV viral replication in Vero E6 cells with an EC_50_ value of 12.9 μM^12^. Several follow up studies have been conducted to optimize the enzymatic inhibition and cellular antiviral activity of this series of compounds, however, no significant improvement has been made^13,14^.

**Fig. 1.**
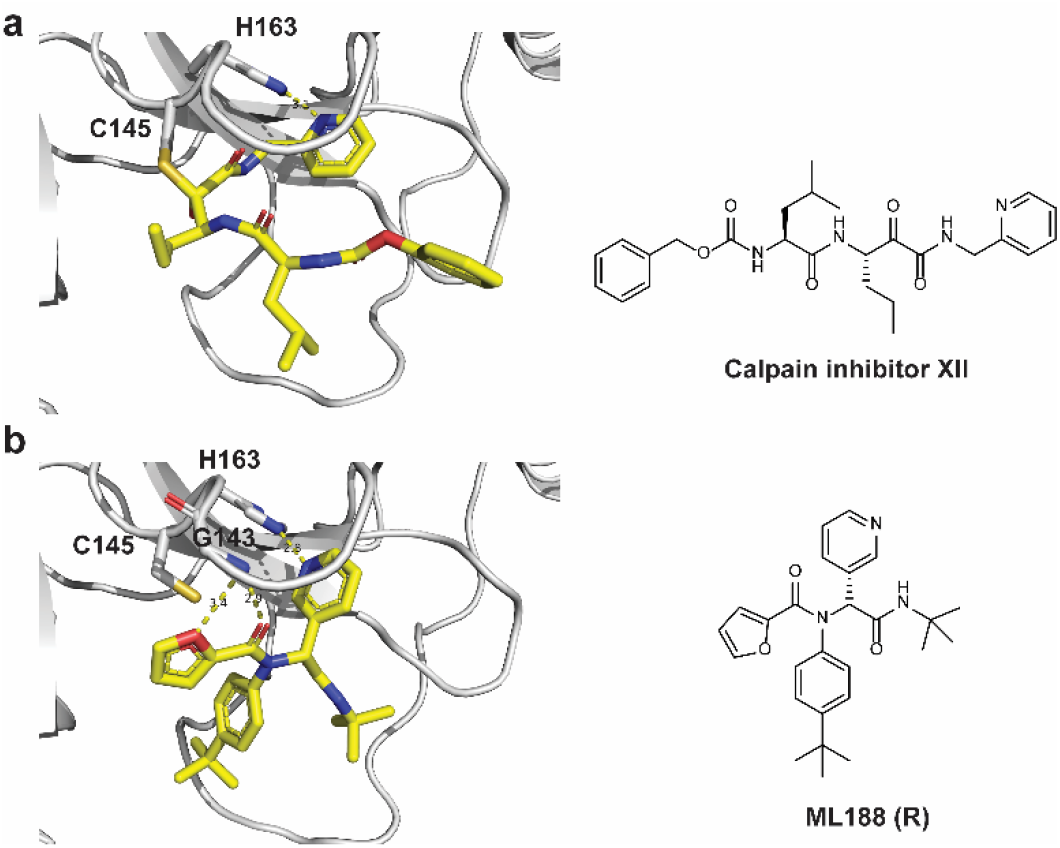
Structure of M^pro^ with inhibitors. **a** X-ray crystal structure of SARS-CoV-2 M^pro^ in complex with calpain inhibitor XII (PDB: 6XFN). **b** X-ray crystal structure of SARS-CoV M^pro^ in complex with ML188 (R) (PDB: 3V3M). Hydrogen bonds are shown in dashed lines.

The similar binding mode of ML188 (R) with calpain inhibitor XII, coupled with the convenient synthesis through the one pot Ugi-4CR, inspired us to design non-covalent SARS-CoV-2 M^pro^ inhibitors based on the ML188 (R) scaffold. Specifically, we leverage our understanding of the M^pro^ inhibition mechanism based on the X-ray co-crystal structures of SARS-CoV-2 M^pro^ with multiple inhibitors to guide the lead optimization (Figs. 2a-d)^7,8^. Overlaying the X-ray crystal structures of SARS-CoV M^pro^ + ML188 (R) (PDB: 3V3M) and the SARS-CoV M^pro^ H41A mutant + the peptide substrate (PDB: 2Q6G) revealed that the furyl, 4-*tert*-butylphenyl, pyridinyl, and *tert*-butyl of ML188 (R) fit in the S1’, S2, S1, and S3 pockets respectively (Figs. 2a and 2d). Therefore, the furyl, 4-*tert*-butylphenyl, pyridinyl, and *tert*-butyl substitutions in ML188 (R) were defined as P1’, P2, P1, and P3, respectively. Next, overlaying the structure of SARS-CoV M^pro^ + ML188 (R) (PDB: 3V3M) and SARS-CoV-2 M^pro^ + GC376 (PDB: 6WTT) suggested that the *tert*-butyl at the P3 substitution of ML188 (R) can be extended to fit in the S4 pocket (Figs. 2b and 2d). Previous structure-activity relationship studies of GC376 indicate that P4 substitution is important, while P3 substitution does not contribute significantly to the binding affinity, as it is solvent exposed^3,8,9,15^. Similarly, the overlaying structures of SARS-CoV M^pro^ + ML188 (R) (PDB: 3V3M) and SARS-CoV-2 M^pro^ + UAWJ247 (PDB: 6XBH) suggested that the 4-*tert*-butyl at the P2 substitution of ML188 (R) can be replaced by phenyl to occupy the extra space in the S2 pocket (Figs. 2c and 2d). Overall, the design mainly focuses on extending the P2 and P3 substitutions of ML188 (R) to achieve optimal shape complementarity with the SARS-CoV-2 M^pro^ (Fig. 2e). In practice, we adopted a stepwise optimization procedure in which the P3 and P2 substitutions were optimized individually in step 1, and then the optimal P2/P3 substitutions were combined in step 2 (Fig. 2e).

**Fig. 2.**
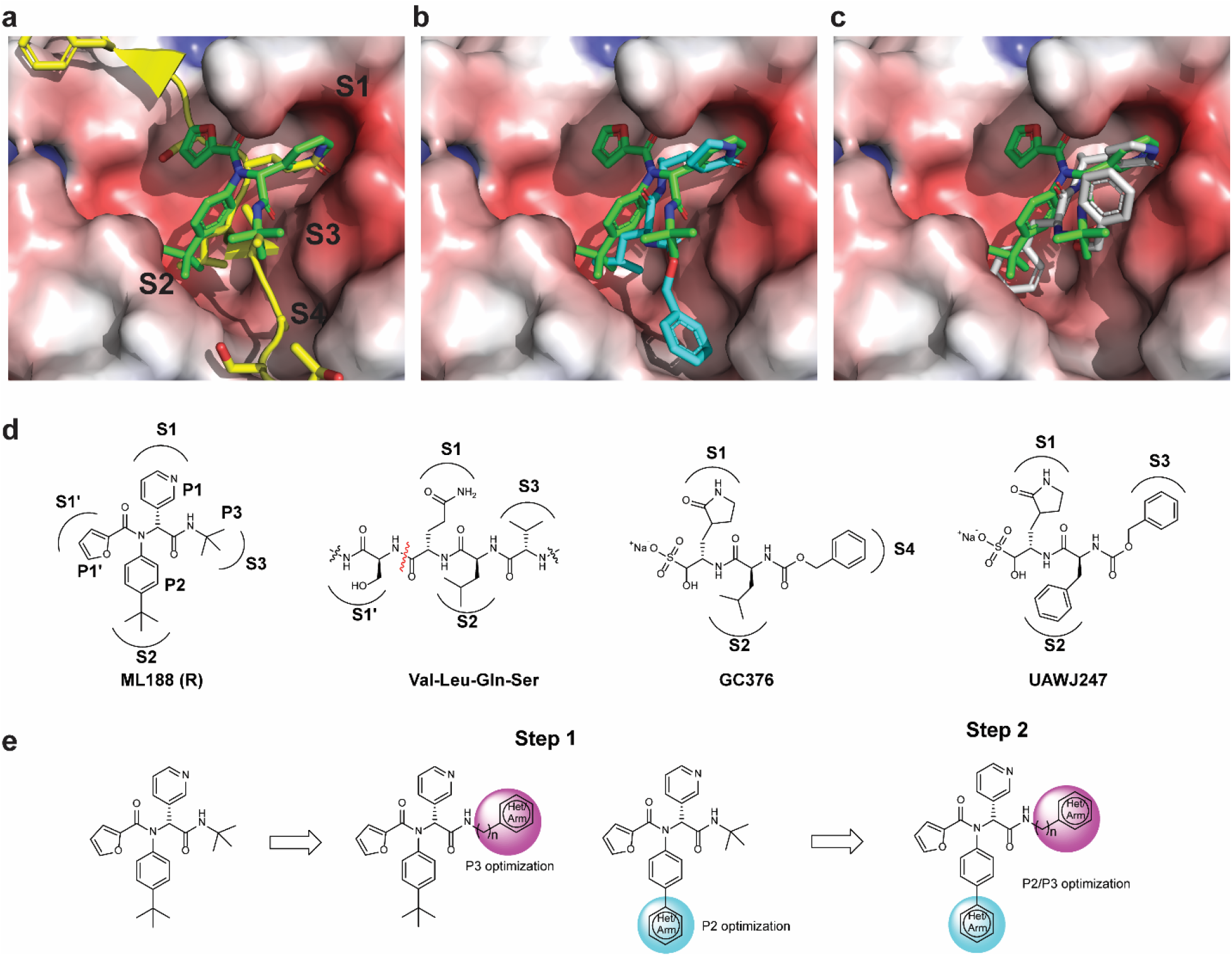
Design rationale for the non-covalent SARS-CoV-2 M^pro^ inhibitors. **a** Overlaying X-ray crystal structures of SARS-CoV M^pro^ + ML188 (R) (PDB: 3V3M, green) and SARS-CoV M^pro^ H41A mutant + peptide substrate (PDB: 2Q6G, yellow with backbone shown as ribbon representation). **b** Overlaying X-ray crystal structures of SARS-CoV M^pro^ + ML188 (R) (PDB: 3V3M) and SARS-CoV-2 M^pro^ + GC376 (PDB: 6WTT). **c** Overlaying X-ray crystal structures of SARS-CoV M^pro^ + ML188 (R) (PDB: 3V3M) and SARS-CoV-2 M^pro^ + UAWJ247 (PDB: 6XBH). **d** Chemical structures of ML188 (R), peptide substrate VLQS, GC376, and UAWJ247. **f** Stepwise optimization of ML188 (R) towards potent non-covalent SARS-CoV-2 M^pro^ inhibitor.

Guided by the design rationale elucidated above, a focused library of ML188 analogs were designed and synthesized (Fig. 3). As the P1’ furyl and P1 pyridinyl both form a critical hydrogen bond with the M^pro^ (Figs. 3a-b), the P1’ and P1 substitutions were kept with minimal variations (Fig. 3c). All designed compounds were synthesized using the one pot Ugi four-component reaction and tested as enantiomer/diastereomer mixtures (Fig. 3c). To circumvent the need of relying on expansive chiral HPLC column for the separation of enantiomers, we strategically introduced the chiral isocyanide so that the diastereomer product mixture can be separated by convenient silica gel column or reverse phase HPLC column purification.

**Fig. 3.**
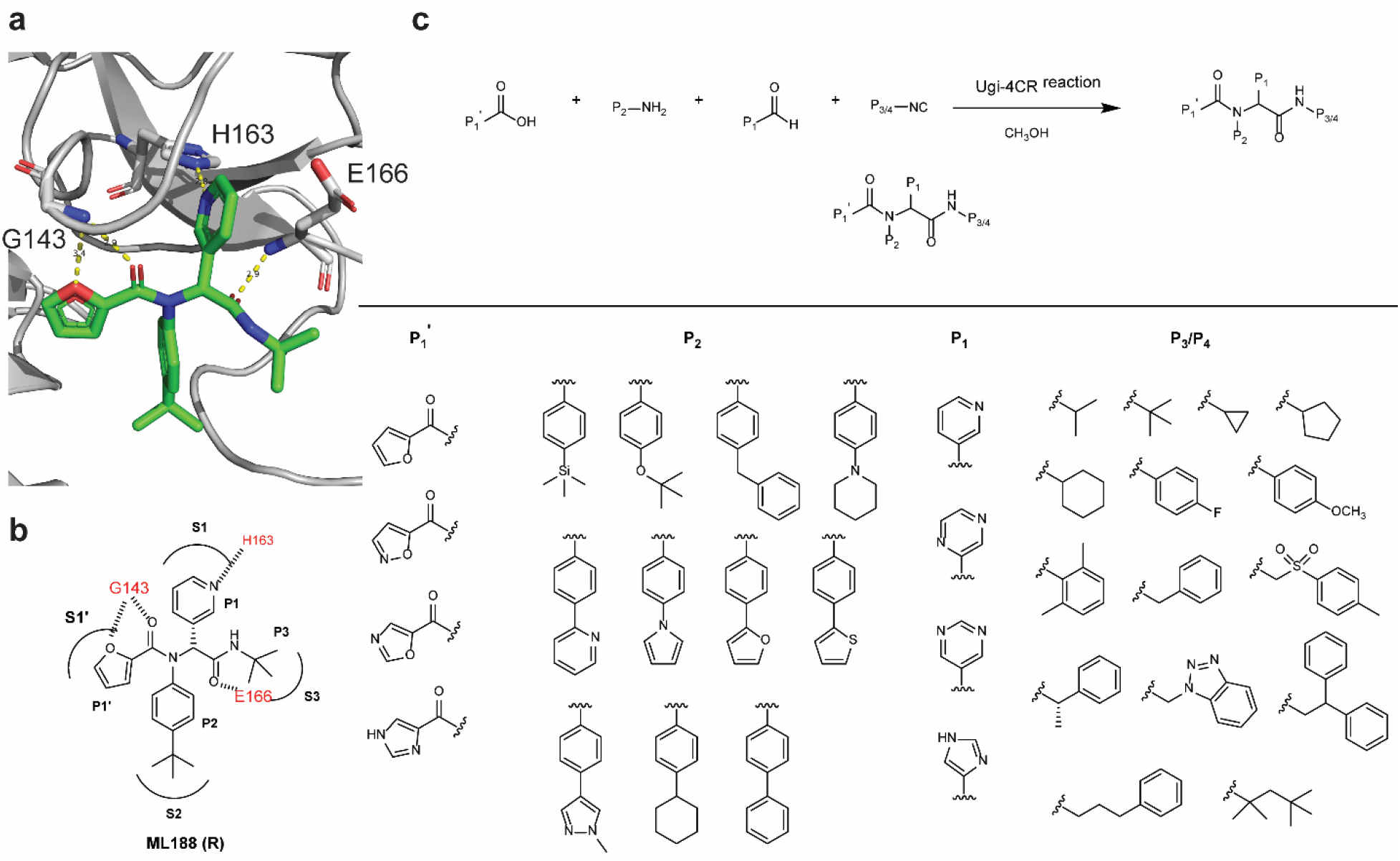
Design and synthesis of a focused library of non-covalent SARS-CoV-2 M^pro^ inhibitors. **a** X-ray crystal structure of SARS-CoV M^pro^ + ML188 (R) (PDB: 3V3M). **b** Binding interactions of ML188 (R) with SARS-CoV M^pro^. **c** Synthesis of ML188 analogs using the Ugi four-component reaction.

### Structure-activity relationship studies of non-covalent SARS-CoV-2 M^pro^ inhibitors

In total, 39 compounds were synthesized (Figs. 4a-4e) and all compounds were initially tested as a mixture of enantiomers or diastereomers in the FRET-based enzymatic assay against SARS-CoV-2 M^pro^ at 20 μM (Fig. 4f). Compounds showing more than 50% inhibition at 20 μM were further titrated to determine the IC_50_ values. Next, compounds with IC_50_ values lower than 5 μM were selected for cellular cytotoxicity profiling in Vero E6 cells, the cell line which was used for the SARS-CoV-2 antiviral assay. The purpose was to prioritize lead candidates for the *in vitro* cellular antiviral assay with infectious SARS-CoV-2. Compounds with potent enzymatic inhibition (IC_50_ < 5 μM) but moderate to high cellular cytotoxicity (CC_50_ < 100 μM) were labeled in red. Compounds with both potent enzymatic inhibition (IC_50_ < 5 μM) and low cellular cytotoxicity (CC_50_ > 100 μM) were labeled in blue (Figs. 4a-4e). As shown in Fig. 4f, the majority of the designed compounds showed more than 50% inhibition when tested at 20 μM. Specifically, Fig. 4a lists compounds with P4 variations. As a reference, ML188 (**1**) (racemic mixture) inhibits SARS-CoV-2 M^pro^ with an IC_50_ value of 10.96 ± 1.58 μM. It was found that compounds **2**, **3**, **5**, **6**, **7**, **8**, **10**, and **13** had improved enzymatic inhibition compared to ML188 (**1**). These results suggest that: a) isopropyl (**2**), cyclopropyl (**3**), cyclopentyl (**5**), cyclohexyl (**6**), and phenyl (**7** and **8**) are the more favorable substitutions at the P3 position than *tert*-butyl; b) compound **13** with the (*S*)-α-methylbenzyl substitution at the P3 position had improved potency, which suggests that extending the substitutions to the S4 pocket might improve the enzymatic inhibition (Fig. 2b). Given the advantage of convenient separation of diastereomers over enantiomers, we therefore decided to fix the P3/P4 substitution as α-methylbenzyl substitution during the P2 optimization (Fig. 4b). All compounds in Fig. 4b were designed to have extended substitutions at the 4-position of benzyl to occupy the extra space in the S2 pocket (Fig. 2c). Consistent with the design hypothesis, several compounds including **14**, **17**, **18**, **19**, **20**, **21**, and **23** had significantly improved enzymatic inhibition (IC_50_ < 3 μM) compared to compound **13**. Replacing the *tert*-butyl in compound **13** with the bulkier trimethylsilyl led to compound **14** with a 2.9-fold increase in M^pro^ inhibition. Cyclohexyl (**17**), thienyl (**19**), pyrrolyl (**20**), pyridinyl (**21**), and phenyl (**23**) were found to be the most favorable substitutions at the S2 pocket. Compound **16** with piperidyl substitution had similar potency as compound **13**, while compound **15** with *O-tert* butyl was less active. Further extending the substitution to benzyl led to compound **22** that was inactive, suggesting biphenyl might be the largest substitution that can be accommodated at the S2 pocket.

**Fig. 4.**
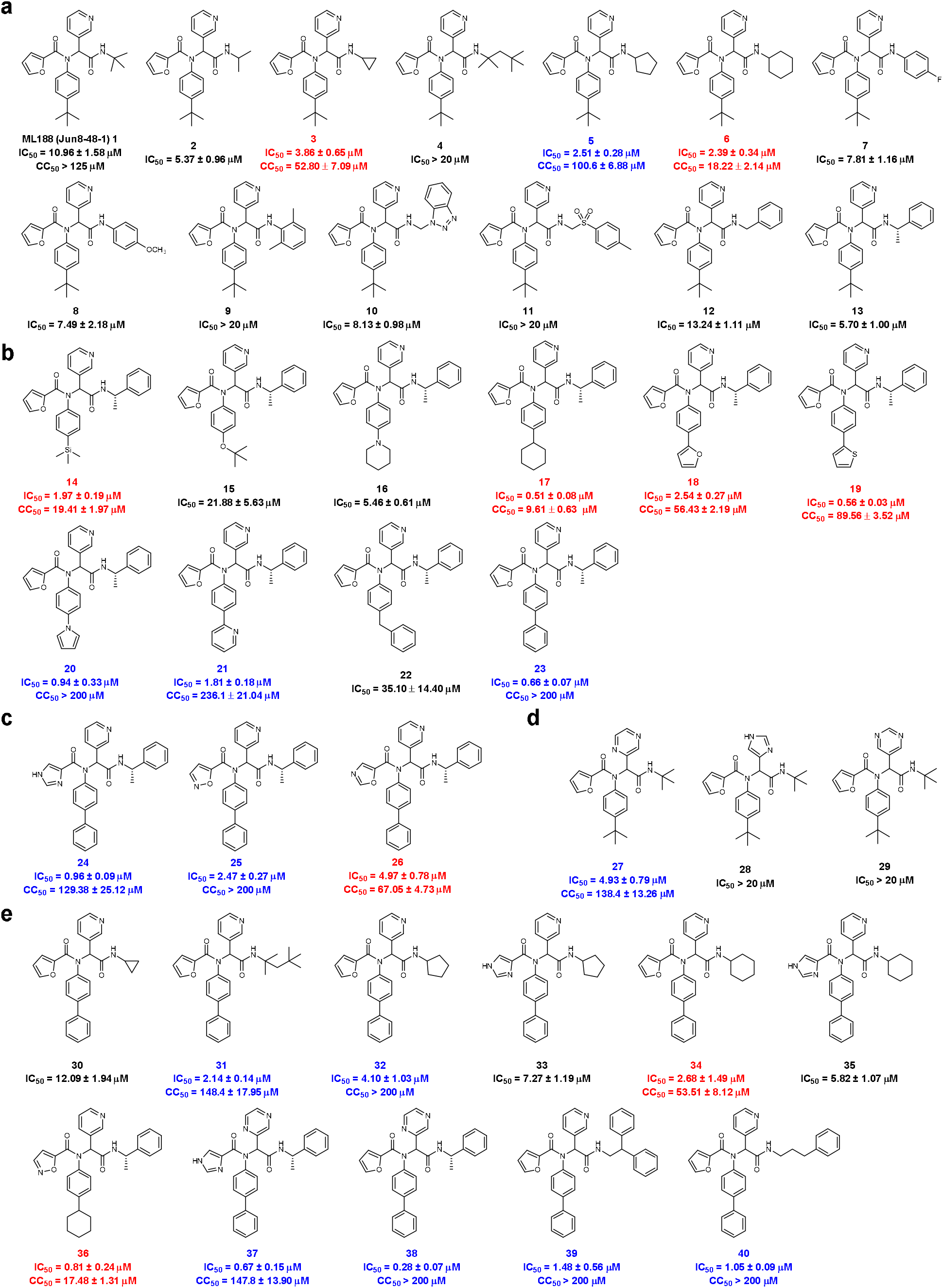
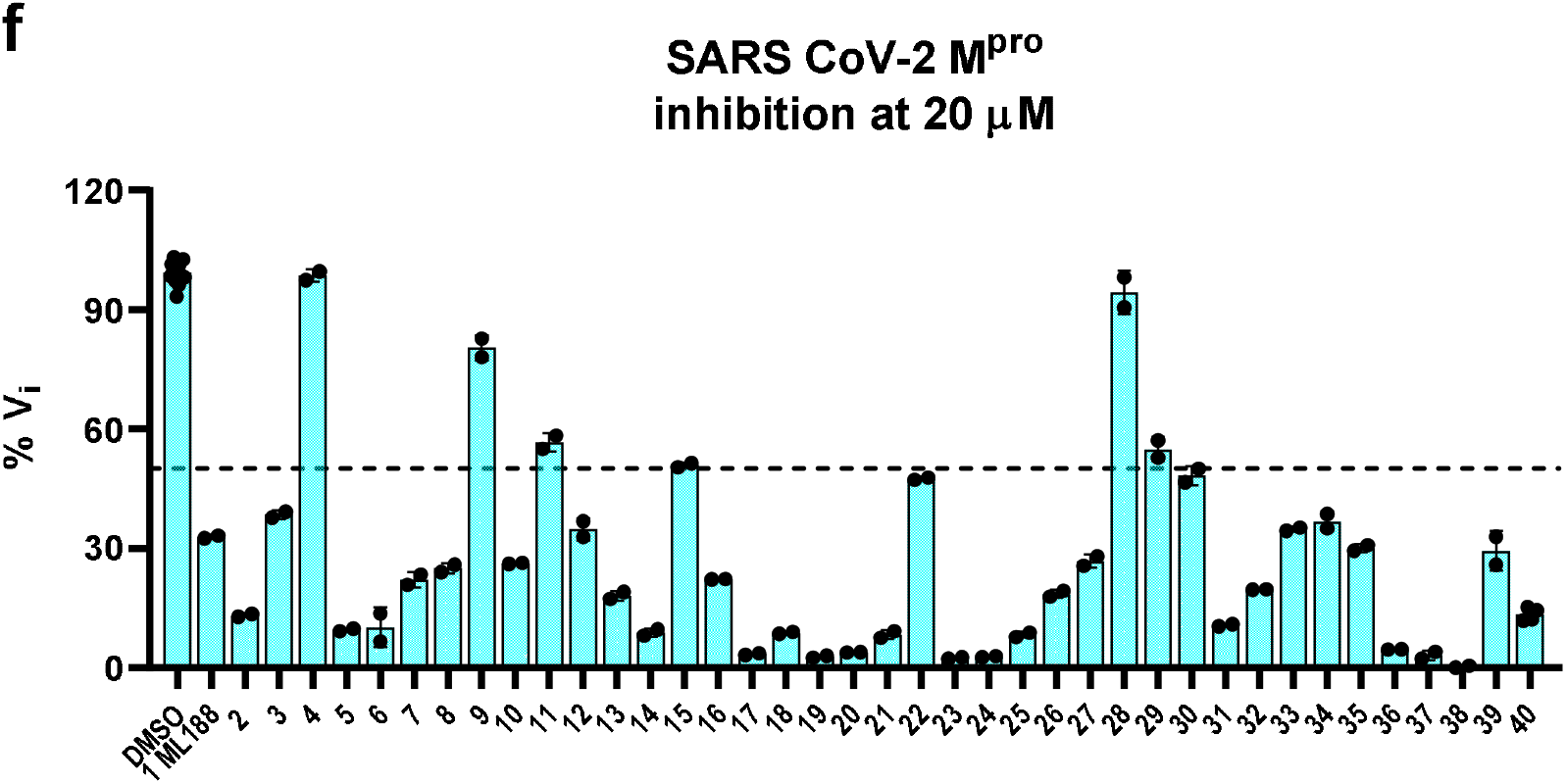
Structures of non-covalent SARS-CoV-2 M^pro^ inhibitors and the enzymatic inhibition against SARS-CoV-2 M^pro^. **a** Analogs with P3/P4 modifications. **b** Analogs with P2 modifications. **c** Analogs with P1’ modifications. **d** Analogs with P1 modifications. **e** Analogs with combined P1’, P1, P2, and P3/P4 modifications. Compounds with potent enzymatic inhibition (IC_50_ < 5 μM) but moderate to high cellular cytotoxicity (CC_50_ < 100 μM) were labeled in red. Compounds with both potent enzymatic inhibition (IC_50_ < 5 μM) and low cellular cytotoxicity (CC_50_ > 100 μM) were labeled in blue. **f** Percentage enzymatic inhibition of SARS-CoV-2 M^pro^ by the designed compounds at 20 μM compound concentration.

The P1’ and P1 substitutions (Figs. 4c and 4d) were chosen to retain the critical hydrogen bonds in ML188 (Fig. 3a). It was found that imidazole (**24**) was tolerated at the P1’ position (IC_50_ = 0.96 ± 0.09 μM), followed by isoxazole (**25**) (IC_50_ = 2.47 ± 0.27 μM), and oxazole (**26**) (IC_50_ = 4.97 ± 0.78 μM). Pyrazine (**27**) was tolerated at the P1 position (IC_50_ = 4.93 ± 0.79 μM); however, pyrimidine (**29**) and imidazole (**30**) were not preferred (IC_50_ > 20 μM).

Next, the above identified favorable P1’, P2, P1, and P3/P4 substitutions were combined and the designed compounds were shown in Fig. 4e. Compounds **36**, **37**, and **38** were the most potent leads with IC_50_ values of 0.81 ± 0.24, 0.67 ± 0.15, and 0.28 ± 0.07 μM, respectively. Compound **39** and **40** were also highly active with IC_50_ values of 1.48 ± 0.56 and 1.05 ± 0.09 μM, respectively.

Among the active compounds with IC_50_ value lower than 5 μM, compounds **3**, **6**, **14**, **17**, **18**, **19**, **26**, **34**, and **36** had moderate to high cellular cytotoxicity in Vero E6 cells (Figs. 4a-4e red), while compounds **5**, **20**, **21**, **23**, **24**, **25**, **27**, **31**, **32**, **37**, **38**, **39**, and **40** were well tolerated and the CC_50_ values were greater than 100 μM.

Next, compounds with potent enzymatic inhibition (IC_50_ ≤ 1 μM) and low cellular cytotoxicity (CC_50_ > 100 μM) were prioritized for the cellular antiviral assay with infectious SARS-CoV-2 in Vero E6 cells using the immunofluorescence assay as the primary assay (Table 1). ML188 (**1**) was included as a control. It was found that ML188 (**1**) was inactive in the antiviral assay (EC_50_ > 20 μM), probably due to its incomplete inhibition of the M^pro^ in the cellular content. Gratifyingly, compounds **20**, **23**, **37**, **38**, and **40** all had potent cellular antiviral activity with EC_50_ values ranging from 0.82 to 4.54 μM. Compound **24** was less active (EC_50_ = 13.06 ± 2.30 μM), possibly due to the poor cellular membrane permeability.

**Table 1.** Antiviral activity and selectivity index of non-covalent SARS-CoV-2 M^pro^ inhibitors. (Selection criteria IC_50_ < 1 μM, CC_50_ > 100 μM).

| Compound ID | SARS-CoV-2 $\text{M}^{\text{pro}}$ $\text{IC}_{50}$ ($\mu\text{M}$) | Vero E6 Cytotoxicity $\text{CC}_{50}$ ($\mu\text{M}$) | SARS-CoV-2 antiviral assay $\text{EC}_{50}$ ($\mu\text{M}$) | Selectivity index SI $\text{CC}_{50}/\text{EC}_{50}$ |
| --- | --- | --- | --- | --- |
| **1 ML188** | $10.96 \pm 1.58$ | $> 125$ | $> 20$ | N.A. |
| **20** | $0.94 \pm 0.33$ | $> 200$ | $1.73 \pm 0.19$ | $> 115.6$ |
| **23** | $0.66 \pm 0.07$ | $> 200$ | $1.61 \pm 0.29$ | $> 124.2$ |
| **24** | $0.96 \pm 0.09$ | $129.38 \pm 25.12$ | $13.06 \pm 2.30$ | 9.9 |
| **37** | $0.67 \pm 0.15$ | $147.8 \pm 13.9$ | $4.54 \pm 0.69$ | 32.6 |
| **38** | $0.28 \pm 0.07$ | $> 200$ | $0.82 \pm 0.56$ | $> 243.9$ |
| **40** | $1.05 \pm 0.09$ | $> 200$ | $2.04 \pm 1.08$ | $> 98.0$ |

Given the potent antiviral activity and a high selectivity index of these potent lead compounds, we then selected compound **23** for further characterization. The two diastereomers of **23** were separated by reverse phase HPLC (Fig. 5). Both diastereomers were tested in the FRET-based enzymatic assay. GC376 was included as a positive control. It was found that **23R** is the active diastereomer with an IC_50_ value of 0.31 ± 0.04 μM, while the **23S** diastereomer was more than 16-fold less active (IC_50_ = 5.61 ± 0.71 μM) (Table 2). The stereochemistry of **23R** was determined by the co-crystal structure with SARS-CoV-2 M^pro^ as described in the following section. Compared with the parent compound ML188 (**1**), the optimized lead **23R** had more than a 35-fold increase in enzymatic inhibition against SARS-CoV-2 M^pro^. Compound **23R** also showed comparable potency against SARS-CoV M^pro^ with an IC_50_ value of 0.27 ± 0.03 μM. Neither ML188 (**1**) nor **23R** inhibited the SARS-CoV-2 papain-like protease (PL^pro^) (IC_50_ > 20 μM) (Table 2), suggesting the inhibition of SARS-CoV-2 M^pro^ by **23R** is specific.

**Fig. 5.**
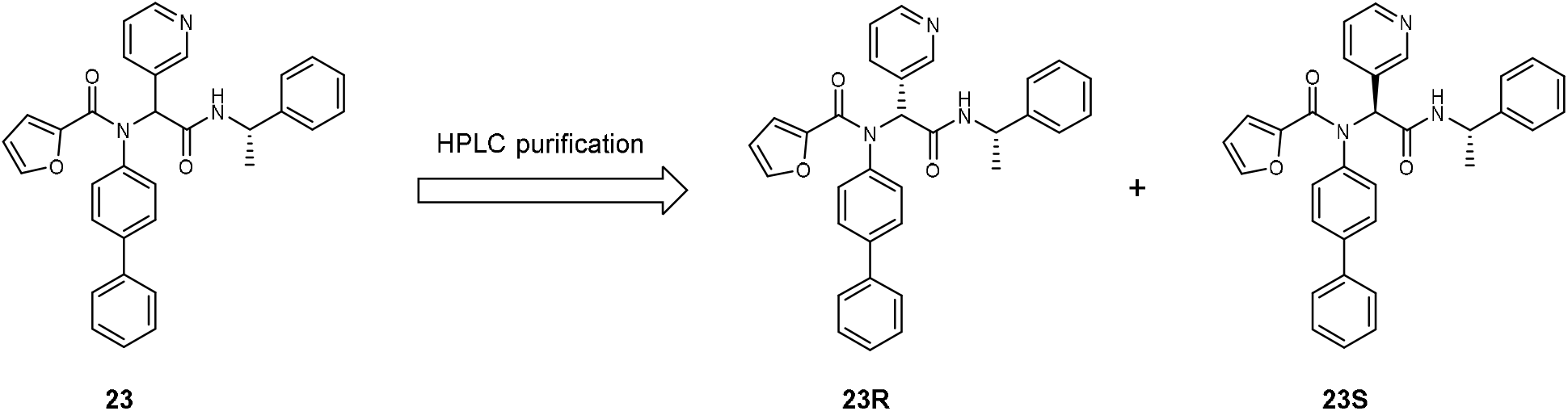
Separation of the two diasteromers of compound 23. The absolute stereochemistry of compound **23R** was determined in the co-crystal structure of this diasteromer with SARS-CoV-2 M^pro^ (PDB: 7KX5).

**Table 2.**
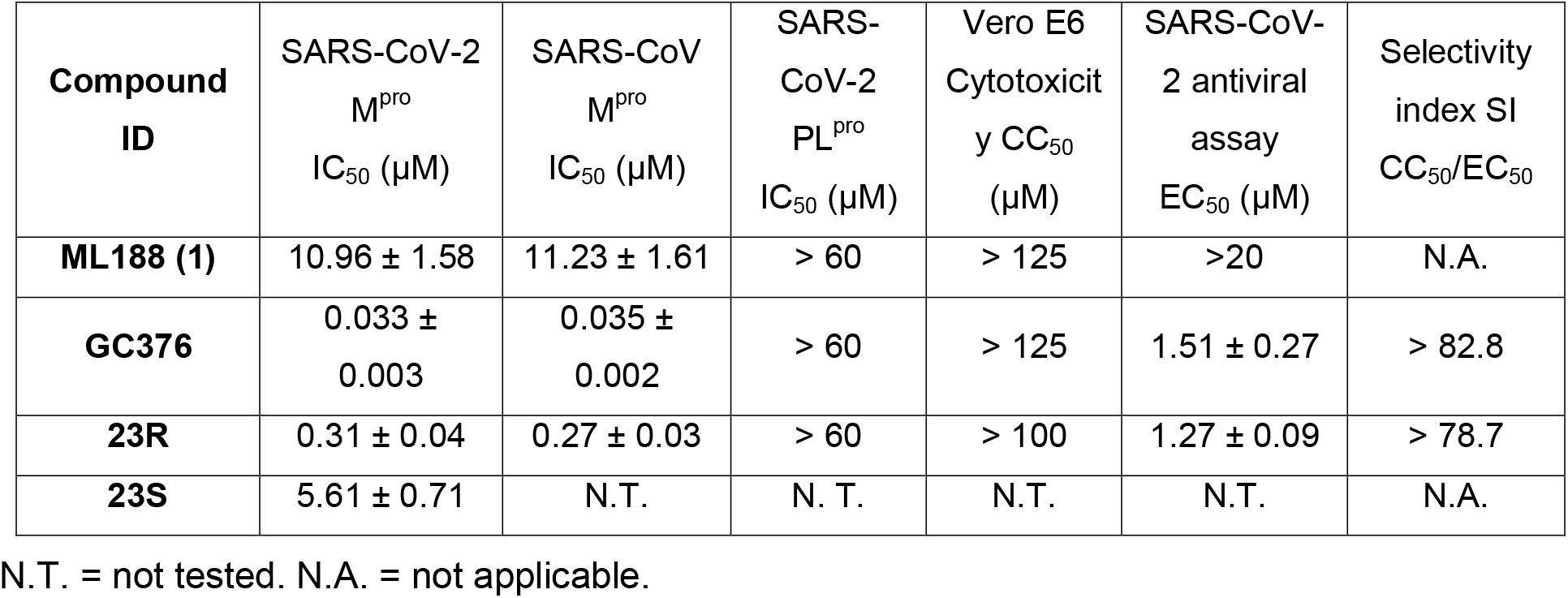
Enzymatic inhibition, antiviral activity and selectivity index of 23R.

Next, the antiviral activity of **23R** was tested against SARS-CoV-2 (USA-WA1/2020 isolate) in Vero E6 cells using the immunofluorescence assay. ML188 (**1**) and GC376 were included as controls. It was found that compound **23R** had an EC_50_ value of 1.27 μM (Table 2), which was similar to the antiviral potency of the covalent inhibitor GC376 (EC_50_ = 1.51 μM). Compound **23R** was also not cytotoxic to Vero E6 cells at up to 100 μM. In contrast, the parent compound ML188 (**1**) had no detectable antiviral activity when tested at up to 20 μM. To further confirm the antiviral activity of compound **23R**, we performed secondary antiviral assay in the human lung epithelial Calu-3 cell line, which endogenously expresses TMPRSS2 and is widely used as a physiological relevant cell line for SARS-CoV-2 infection. It was found that compound **23R** inhibited SARS-CoV-2 (USA-WA1/2020 isolate) replication in Calu-3 cells with an EC_50_ value of 3.03 μM and it was not cytotoxic at up to 100 μM (Fig. 6). The 2.4-fold difference in antiviral potency between the Vero E6 and Calu-3 cell lines might due to differences in cell membrane permeability or metabolism.

**Fig. 6.**
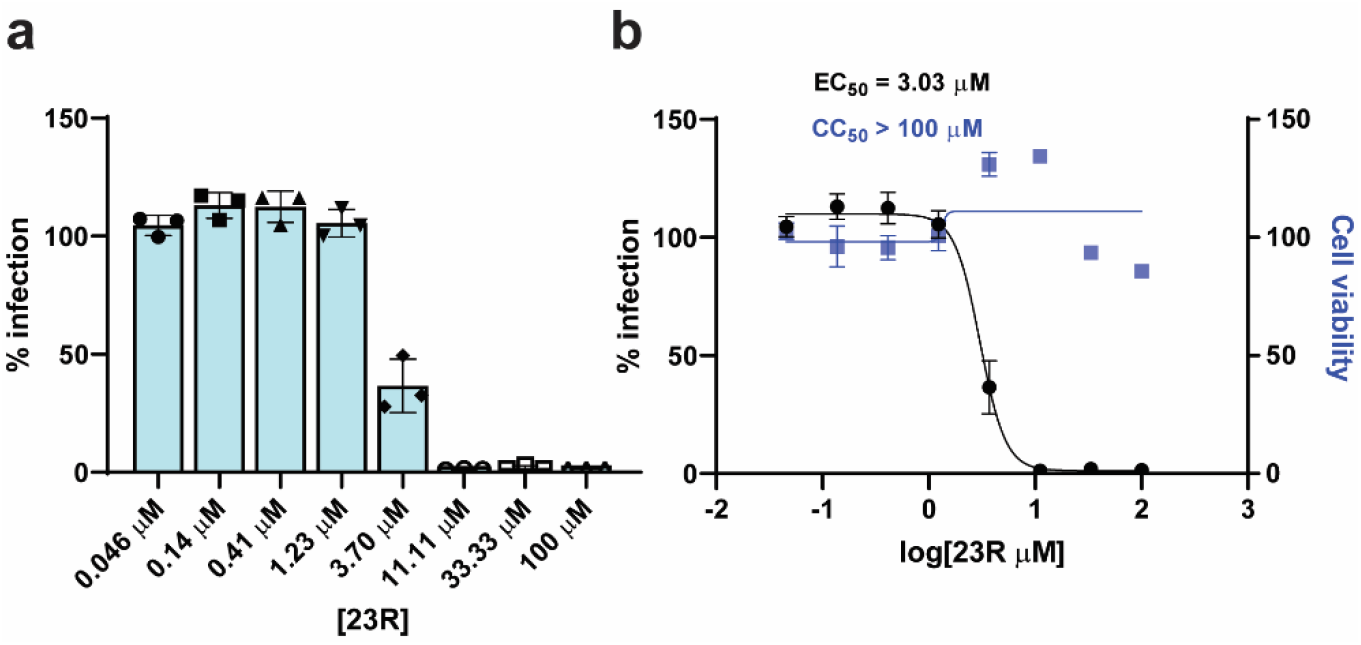
Antiviral activity of 23R against SARS-CoV-2 in Calu-3 cells. **a** Raw data of the percentage of immunofluorescence positive cells with different concentrations of **23R**. **b** Antiviral potency and cytotoxicity plots.

### Mechanism of action of 23R in inhibiting SARS-CoV-2 M^pro^

The mechanism of action of **23R** was characterized using the native mass spectrometry binding assay, the thermal shift binding assay, and the enzymatic kinetic studies (Fig. 7). In the native mass spectrometry binding assay, compound **23R** showed dose-dependent binding to SARS-CoV-2 M^pro^, similar to the positive control GC376, with a binding stoichiometry of one drug per monomer (Fig. 7a). Similarly, compound **23R** showed dose-dependent stabilization of the SARS-CoV-2 M^pro^ in the thermal shift binding assay with an apparent Kd value of 9.43 μM, a 9.3-fold increase compared to ML188 (**1**) (Fig. 7b). In the enzymatic kinetic studies, **23R** was shown to be a reversible inhibitor with a Ki value of 0.07 μM (Figs. 7c and 7d top and middle panels). In comparison, the Ki for the parent compound ML188 (**1**) is 2.29 μM. The Lineweaver-Burk or double-reciprocal plot with different compound concentrations yielded an intercept at the Y-axis, suggesting that **23R** is a competitive inhibitor similar to ML188 (**1**) (Figs. 7c and 7d bottom panel).

**Fig. 7.**
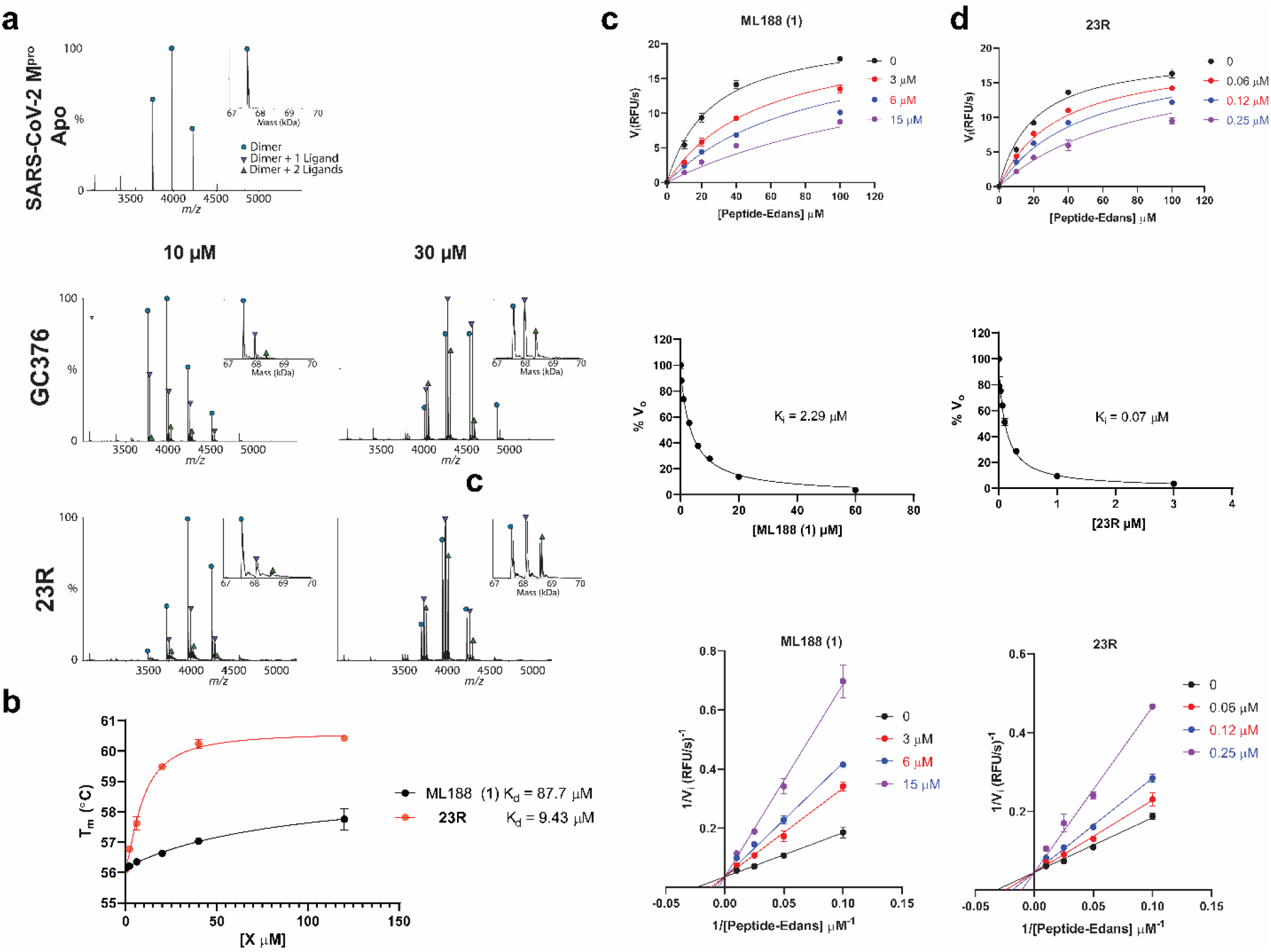
Characterization of the mechanism of action of 23R to SARS-CoV-2 M^pro^ using native mass-spectrometry, thermal shift assay, and enzyme kinetic studies. **a** Binding of **23R** to SARS-CoV-2 M^pro^ using native mass spectrometry. Native mass spectra with the inset deconvolved spectra revealing ligand binding with 10 μM or 30 μM GC-376 added (middle panel), and 10 μM and 30 μM **23R** (bottom panel) with 4 mM DTT added. The peaks are annotated with the blue circle as the dimer, green down triangle as the dimer with one ligand bound, and the purple up triangle as the dimer with two ligands bound. **b** Dose-dependent melting temperature (Tm) shift in thermal shift assay. 3 μM SARS-CoV-2 M^pro^ protein was incubated with various concentrations of ML188 or **23R** in the presence of 4 mM DTT. Measured T_m_ was plotted against compound concentration with one-site binding function in Prism 8. **c**, **d** Enzymatic kinetic assay with ML188 and compound **23R**. Kinetic parameters in the presence of various concentrations of ML188 or **23R** were globally fit with Michalis-Menten function in prism 8 (top panels); double reciprocal plots were shown in the bottom panels. The middle panels show the Morrison plots of compound ML188 and **23R** with 20 μM FRET substrate was used.

### X-ray crystal structure of SARS-CoV-2 M^pro^ with 23R

Using X-ray crystallography, we successfully determined the binding pose of **23R** with SARS-CoV-2 M^pro^ at 2.6 Å resolution (Fig. 8a). Electron density reveals the body of **23R** extends throughout the substrate binding channel, with side chains occupying the S1’, S1, S2, and S3 sub-pockets. The binding pose is similar to the previously solved structure of SARS-CoV M^pro^ with ML188 (R) (PDB: 3V3M)^12^, consistent with the similarities between the two compounds and between the two proteins (Fig. 8b). The furyl moiety of **23R** binds to a portion of the P1’ site, which normally accommodates small hydrophobic residues. While the furylamide carbonyl group of **23R** does not insert into the oxyanion hole, it does form a bifurcated hydrogen bond with the apical residue of this oxyanion hole, Gly143. However, the furan ring oxygen is likely a weaker hydrogen bond acceptor than the amide oxygen and it lies outside of the plane of Gly143’s amide NH. Directly attached to the furylamide moiety is a P2 biphenyl group and a P1 pyridinyl ring. The P2 biphenyl group projects directly into the S2 pocket, which prefers hydrophobic residues such as leucine and phenylalanine. As expected, the P1 pyridinyl ring occupies the S1 pocket, which is known for its strict preference for glutamine. While most M^pro^ inhibitors bear a pyrrolidinone glutamine mimetic at the P1 position, we determined that more hydrophobic residues can also bind to the S1 site, and that hydrogen bond formation with His163 is critical for inhibition^8^. In this instance, the pyridinyl ring of **23R** is nearly superimposable with the same moiety from calpain inhibitor XII (Fig. 8c) forming a close (2.9 Å) hydrogen bond with His163. An amide bond connecting the pyridinyl ring to the α-methylbenzyl moiety forms a hydrogen bond with the main chain of Glu166. The benzyl ring of the α-methylbenzyl moiety is partially positioned both in the S2 and S3 pockets, a novel binding pose that has not been observed with existing M^pro^ inhibitors. Normally, a substituent at this position would be expected to flip away from the enzyme core towards the solvent-exposed S3 pocket, which explains why P3 substitutions have little to no influence on the enzymatic inhibition^4^. However, the hydrophobic nature of the benzyl ring in **23R** causes it to project towards the core near the S2 pocket, forcing Gln189 to rotate outwards (Fig. 8d). This conformation is reinforced by pi-stacking interactions with the first phenyl of the biphenyl substituent. Notably, the binding pose of **23R** features continuous intramolecular pi-stacking, where the phenyl is sandwiched by furan and benzyl groups, potentially contributing to its potent inhibition of M^pro^. Meanwhile, the S4 pocket remains largely unoccupied by **23R**, leaving room for further improvement. In summary, the X-ray crystal structure of SARS-CoV-2 M^pro^ in complex with **23R** revealed two interesting structural features: 1) The P2 biphenyl is probably the largest substitution that can be accommodated in the S2 pocket, which is consistent with our design hypothesis. 2) The benzyl group from the terminal α-methylbenzyl fits in a previously unexplored binding pocket located in between the S2 and S4 pockets. As such, the benzyl group faces towards the core of the enzyme instead of solvent-exposed as seen with other existing M^pro^ inhibitors. Although this is unexpected from the design perspective, this novel binding mode suggests that the new binding pocket in between S2 and S4 that can be explored for inhibitor design.

**Fig. 8.**
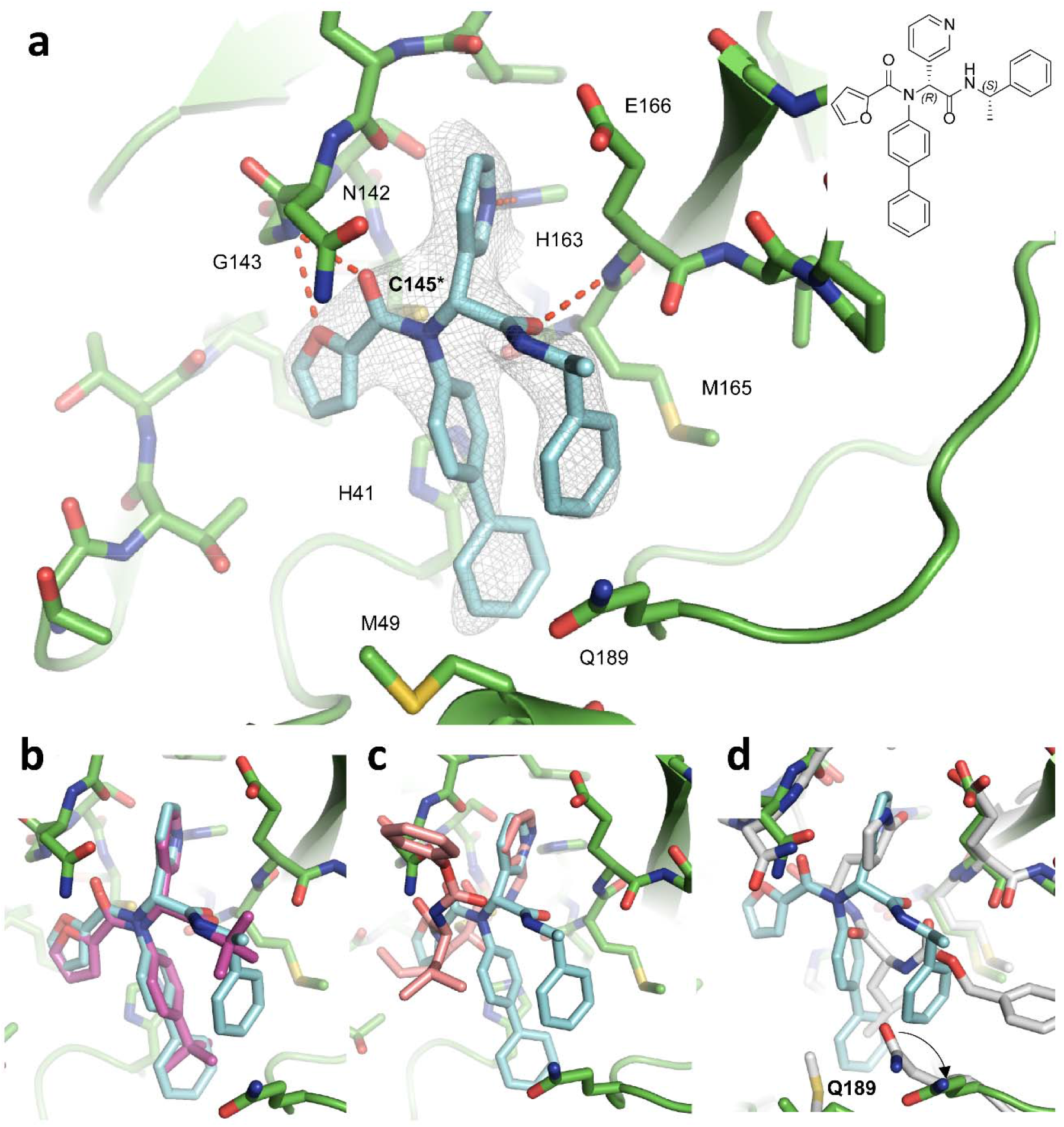
X-ray crystal structure of SARS-CoV-2 M^pro^ in complex with 23R. **a** X-ray crystal structure of SARS-CoV-2 M^pro^ in complex with **23R** (PDB: 7KX5). **b** Overlaying structures of SARS-CoV-2 M^pro^ + **23R** (PDB: 7KX5) and SARS-CoV M^pro^ + ML188 (R) (PDB: 3V3M). **c** Overlaying structures of SARS-CoV-2 M^pro^ + **23R** (PDB: 7KX5) and SARS-CoV-2 M^pro^ + calpain inhibitor XII (PDB: 6XFN). **d** Overlaying structures of SARS-CoV-2 M^pro^ + **23R** (PDB: 7KX5) and SARS-CoV-2 M^pro^ + GC376 (PDB: 6WTT).

## Discussion

Given the social and economic impact of the ongoing COVID-19 pandemic, the need for effective therapeutics is apparent. Researchers around the globe are racing to come up with intervention strategies. The viral M^pro^ is a high profile antiviral drug target and several M^pro^ inhibitors are now in animal model studies and human clinical trial^6^. Among the known M^pro^ inhibitors, the majority of them are covalent inhibitors such as GC376 analogs that contain a pyrollidone in the P1 position as a glutamine mimetic. Several structurally distinct compounds including ebselen, disulfiram, carmofur, PX-12, tideglusib, and shiknonin were claimed as M^pro^ inhibitors^16,17^, but were later invalidated as promiscuous non-specific cysteine protease inhibitors^18,19^. In addition, non-covalent inhibitors such as ML188 (R) were developed and validated as SARS-CoV M^pro^ inhibitors^5,12^. Several follow up studies have been conducted to optimize the enzymatic potency of this series of compounds against SARS-CoV M^pro^ and the SARS-CoV-2 M^pro^. However, no significant improvement has been made and ML188 (R) remains the only non-covalent inhibitor with moderate antiviral activity against SARS-CoV (EC_50_ = 12.9 ± 0.7 μM)^13,14^. Nevertheless, given the sequence and structure similarities between SARS-CoV and SARS-CoV-2 M^pro^, and the common binding mode of ML188 (R) and calpain inhibitor XII, we hypothesize that ML188 (R) is a promising scaffold for lead optimization. In this study, we developed an expedited drug discovery approach to the design of non-covalent inhibitors of SARS-CoV-2 M^pro^. The design was based on the binding pose of calpain inhibitor XII with SARS-CoV-2 M^pro^ (PDB: 6XFN). The highlights of this study include: 1) The overlaying X-ray crystal structures with multiple M^pro^ inhibitors revealed the chemical space that can be explored for drug design. 2) All designed compounds were synthesized by the one-pot Ugi-4CR, which greatly facilitated the lead optimization. Indeed, we were able to improve the enzymatic inhibition potency by 35-fold from a focused library of 39 compounds. This is a significant advantage compared to covalent inhibitors such as GC376, which involves at least a five-step synthesis. 3) By introducing the chiral isocyanide, the diastereomer product can be conveniently separated by either silica gel column or reverse phase HPLC column, bypassing the need for an expensive chiral HPLC column. This greatly speeds up the co-crystallization. 4) The X-ray crystal structure of SARS-CoV-2 M^pro^ in complex with **23R** reveals a new binding pocket in between S2 and S4 sites that can be explored for drug design. Overall, using the expedited drug discovery approach, this study revealed a promising non-covalent M^pro^ inhibitor **23R** with a confirmed mechanism of action and potent cellular antiviral activity for further development.

## MATERIALS

### Cell lines and viruses

VERO E6 cells (ATCC, CRL-1586) were cultured in Dulbecco’s modified Eagle’s medium (DMEM), supplemented with 5% heat inactivated FBS in a 37°C incubator with 5% CO_2_. SARS-CoV-2, isolate USA-WA1/2020 (NR-52281), was obtained through BEI Resources and propagated once on VERO E6 cells before it was used for this study. Studies involving the SARS-CoV-2 were performed at the UTHSCSA biosafety level-3 laboratory by personnel wearing powered air purifying respirators.

### Protein expression and purification

SARS CoV-2 main protease (M^pro^ or 3CL) gene from strain BetaCoV/Wuhan/WIV04/2019 and SARS-CoV main protease from strain CDC#200301157 in the pET29a(+) vector with *E. coli* codon optimization were ordered from GenScript (Piscataway, NJ). The M^pro^ gene was then subcloned into pE-SUMO vector as described previously^7,8^. The expression and purification of SARS-CoV and SARS-CoV-2 M^pro^ with unmodified N- and C-termini was detailed in our previous publication^8^.

The expression and purification of SARS CoV-2 papain-like protease (PL^pro^) was also described in our previous publications^7,8,18^.

### Peptide synthesis

The SARS-CoV-2 M^pro^ FRET substrate Dabcyl-KTSAVLQ/SGFRKME(Edans) was synthesized as described before.^7^ The SARS-CoV-2 PL^pro^ FRET substrate Dabcyl-FTLRGG/APTKV(Edans) was synthesized by solid-phase synthesis through iterative cycles of coupling and deprotection using the previously optimized procedure.^20^

### Compound synthesis and characterization

Details for the synthesis procedure and characterization for compounds can be found in the supplementary information.

### Native Mass Spectrometry

Prior to analysis, the protein was buffer exchanged into 0.2 M ammonium acetate (pH 6.8) and diluted to 10 μM. DTT was dissolved in water and prepared at a 400 mM stock. Each ligand was dissolved in ethanol and diluted to 10X stock concentrations. The final mixture was prepared by adding 4 μL protein, 0.5 μL DTT stock, and 0.5 μL ligand stock for final concentration of 4 mM DTT and 70 μM protein. Final ligand concentrations were 10 μM and 30 μM. The mixtures were then incubated for 10 minutes at room temperature prior to analysis. Each sample was mixed and analyzed in triplicate.

Native mass spectrometry (MS) was performed using a Q-Exactive HF quadrupole-Orbitrap mass spectrometer with the Ultra-High Mass Range research modifications (Thermo Fisher Scientific). Samples were ionized using nano-electrospray ionization in positive ion mode using 1.0 kV capillary voltage at a 150 °C capillary temperature. The samples were all analyzed with a 1,000–25,000 m/z range, the resolution set to 30,000, and a trapping gas pressure set to 3. Between 10 and 50 V of source fragmentation was applied to all samples to aid in desolvation. Data were deconvolved and analyzed with UniDec.^21^

### Enzymatic assays

The main protease (M^pro^) enzymatic assays were carried out in M^pro^ reaction buffer containing 20 mM HEPES pH 6.5, 120 mM NaCl, 0.4 mM EDTA, 20% glycerol and 4 mM DTT, and the SARS-CoV-2 papain-like protease (PL^pro^) enzymatic assays were carried out in PL^Pro^ reaction buffer containing 50 mM HEPES, pH7.5, 0.01% triton-100 and 5 mM DTT. The percentage of inhibition and enzymatic IC_50_ values were calculated as previously described^7,8^. Briefly, the assay was performed in 96-well plates with 100 μl of 100 nM M^pro^ protein or 200 nM PL^Pro^ protein in their respective reaction buffers. Then 1 μl testing compound at various concentrations was added to each well and incubated at 30 °C for 30 min. The enzymatic reaction was initiated by adding 1 μl of 1 mM corresponding FRET substrate (the final substrate concentration is 10 μM). The reaction was monitored in a Cytation 5 image reader with filters for excitation at 360/40 nm and emission at 460/40 nm at 30 °C for 1 hr. The initial velocity of the enzymatic reaction with and without testing compounds was calculated by linear regression for the first 15 min of the kinetic progress curve.

For the Morrison plot, 10 μl 100 nM SARS-CoV-2 M^pro^ protein was added to 190 μl of M^pro^ reaction buffer containing testing compound and the FRET substrate, and the reaction was monitored for 2 hr. The final FRET substrate concentration in this assay is 20 μM. KI was determined with Morrison equation in Prism 8 as described previously^7,8^.

For Michealis-Menten and Lineweaver-Burk plots, assay was carried as follows: 50 μl of 50 μM M^pro^ protein was added to 50 μl reaction buffer containing testing compound and various concentrations of FRET substrate to initiate the enzyme reaction. The initial velocity of the enzymatic reaction with and without testing compounds was calculated by linear regression for the first 15 min of the kinetic progress curve, the plotted against substrate concentrations in Prism 8 with Michaelis-Menten equation.

### Differential scanning fluorimetry (DSF)

The thermal shift binding assay (TSA) was carried out using a Thermal Fisher QuantStudio™ 5 Real-Time PCR System as described previously^7,8^. Briefly, 3 μM SARS-CoV-2 M^pro^ protein in M^pro^ reaction buffer was incubated with various concentrations of compound ML188 or **23R** at 30 °C for 30 min. 1X SYPRO orange dye was added and fluorescence of the well was monitored under a temperature gradient range from 20 °C to 90 °C with 0.05 °C/s incremental step. Measured T_m_ was plotted against compound concentration with one-site binding function in Prism 8.

### Cytotoxicity measurement

Evaluation of the cytotoxicity of compounds were carried out using the neutral red uptake assay^22,23^. Briefly, 80,000 cells/mL of the tested cell lines were dispensed into 96-well cell culture plates at 100 μL/well. Twenty-four hours later, the growth medium was removed and washed with 150 μL PBS buffer. 200 μL fresh serum-free medium containing serial diluted compounds was added to each well. After incubating for 5 days at 37 °C, the medium was removed and replaced with 100 μL DMEM medium containing 40 μg/mL neutral red and incubated for 2-4 h at 37 °C. The amount of neutral red taken up was determined by measuring the absorbance at 540 nm using a Multiskan FC Microplate Photometer (Fisher Scientific). The CC_50_ values were calculated from best-fit dose response curves with variable slope in Prism 8.

### Immunofluorescence assay

Antiviral immunofluorescence assay was carried out as previously described^8^. Briefly, Vero E6 cells in 96-well plates (Corning) were infected with SARS-CoV-2 (USA-WA1/2020 isolate) at MOI of 0.05 in DMEM supplemented with 1% FBS. Immediately before the viral inoculation, the tested compounds in a three-fold dilution concentration series were also added to the wells in triplicate. The infection proceeded for 48 h without the removal of the viruses or the compounds. The cells were then fixed with 4% paraformaldehyde, permeabilized with 0.1% Triton-100, blocked with DMEM containing 10% FBS, and stained with a rabbit monoclonal antibody against SARS-CoV-2 NP (GeneTex, GTX635679) and an Alexa Fluor 488-conjugated goat anti-mouse secondary antibody (ThermoFisher Scientific). Hoechst 33342 was added in the final step to counterstain the nuclei. Fluorescence images of approximately ten thousand cells were acquired per well with a 10x objective in a Cytation 5 (BioTek). The total number of cells, as indicated by the nuclei staining, and the fraction of the infected cells, as indicated by the NP staining, were quantified with the cellular analysis module of the Gen5 software (BioTek).

### Antiviral assay in Calu-3 cells

Calu-3 cells (ATCC, HTB-55) grown in Minimal Eagles Medium supplemented with 0.1% non-essential amino acids, 0.1% penicillin/streptomycin, and 10% FBS are plated in 384 well plates. The next day, 50 nL of drug suspended in DMSO is added as an 8-pt dose response with three-fold dilutions between test concentrations in triplicate, starting at 40 μM final concentration. The negative control (DMSO, n=32) and positive control (10 μM Remdesivir, n=32) are included on each assay plate. Calu3 cells are pretreated with controls and test drugs (in triplicate) for 2 hours prior to infection. In BSL3 containment, SARS-CoV-2 (isolate USA WA1/2020) diluted in serum free growth medium is added to plates to achieve an MOI=0.5. Cells are incubated continuously with drugs and SARS-CoV-2 for 48 hours. Cells are fixed and then immunstained with anti-dsRNA (J2) and nuclei are counterstained with Hoechst 33342 for automated microscopy. Automated image analysis quantifies the number of cells per well (toxicity) and the percentage of infected cells (dsRNA+ cells/cell number) per well. SARS-CoV-2 infection at each drug concentration was normalized to aggregated DMSO plate control wells and expressed as percentage-of-control (POC=% Infection _sample_/Avg % Infection _DMSO cont_). A non-linear regression curve fit analysis (GraphPad Prism 8) of POC Infection and cell viability versus the log_10_ transformed concentration values to calculate IC_50_ values for Infection and CC_50_ values for cell viability. Selectivity index (SI) was calculated as a ratio of drug’s CC_50_ and IC_50_ values (SI = CC_50_/IC_50_).

### M^pro^ crystallization and structure determination

**23R** was added to 20 mg/mL SARS-CoV-2 M^pro^ to a final concentration of 1.75 mM and incubated overnight at 4°C. This mixture was then diluted four-fold with protein stock buffer (20 mM Tris pH 7.5, 200 mM NaCl, 1 mM DTT) then spun down at 13,000 × g for 1 min to remove precipitate. Crystals were grown by mixing the protein-inhibitor sample with an equal volume of crystallization buffer (20% PEG 3350, 0.2 M NaF) in a vapor diffusion, hanging drop apparatus. Crystals were then transferred to a drop with crystallization buffer containing 5 mM **23R** for 1 h, followed by a brief soaking in a cryoprotectant solution of 30% PEG 3350 and 15% glycerol with 2 mM **23R**. Crystals were then flash frozen in liquid nitrogen for X-ray diffraction.

X-ray diffraction data for the SARS-CoV-2 M^pro^ structures were collected on the SBC 19-ID beamline at the Advanced Photon Source (APS) in Argonne, IL, and processed with the HKL3000 software suite. The CCP4 versions of MOLREP^24^ was used for molecular replacement using a previously solved SARS-CoV-2 M^pro^ structure, 6YB7. Structural refinement was performed using REFMAC5^25^ and COOT^26^. The crystallographic statistics is shown in Table S1.

## Data availability

The complex structure for SARS-CoV-2 M^pro^ with **23R** has been deposited in the Protein Data Bank with the accession ID of 7KX5 (SARS-CoV-2 M^Pro^ + Jun8-76-3A).

## Acknowledgements

This research was partially supported by the National Institutes of Health (NIH) (Grants AI147325 and AI157046) and the Arizona Biomedical Research Centre Young Investigator grant (ADHS18-198859) to J. W. The antiviral assay in Calu-3 cells was conducted through the NIAID preclinical service under a non-clinical evaluation agreement. J. A. T. and M. T. M. were funded by the National Institute of General Medical Sciences and National Institutes of Health (Grant R35 GM128624 to M. T. M.). We thank Michael Kemp for assistance with crystallization and X-ray diffraction data collection. We also thank the staff members of the Advanced Photon Source of Argonne National Laboratory, particularly those at the Structural Biology Center (SBC), with X-ray diffraction data collection. SBC-CAT is operated by UChicago Argonne, LLC, for the U.S. Department of Energy, Office of Biological and Environmental Research under contract DE-AC02-06CH11357. The SARS-CoV-2 experiments were supported by a COVID-19 pilot grant from UTHSCSA and NIH grant AI151638 to Y.X. SARS-Related Coronavirus 2, Isolate USA-WA1/2020 (NR-52281) was deposited by the Centers for Disease Control and Prevention and obtained through BEI Resources, NIAID, NIH.

## AUTHOR CONTRIBUTIONS

J. W. conceived and designed the study; N. K. synthesized and characterized the compounds; M. S. carried out M^pro^ crystallization and structure determination with the assistance of X. Z, and analyzed the data with Y. C.; C. M. expressed the M^pro^ and PL^pro^, and performed the IC_50_ determination, thermal shift-binding assay, and enzymatic kinetic studies; Y. H. performed the cytotoxicity assay; X. M. and F. Z. performed the SARS-CoV-2 immunofluorescence assay in Vero E6 cells under the guidance of Y. X. D. S. performed the SARS-CoV-2 immunofluorescence assay in Calu-3 cells under the guidance of S. C. J. T. performed the native mass spectrometry experiments with the guidance from M. T. M.; J. W. and Y. C. secured funding and supervised the study; J. W., Y.C., and M.S. wrote the manuscript with the input from others.

## Additional Information

Supplementary information accompanies this paper at

