## Supplementary Information for "An expedited approach towards the rationale design of non-covalent SARS-CoV-2 main protease inhibitors with in vitro antiviral activity"

### SUPPLEMENTARY TABLE

### Table S1. Crystallographic statistics

| **Data Collection** | **PDB ID 7KX5** |
|---|---|
| Inhibitor | Jun8-76-3A (**23R**) |
| Space Group | C 1 2 1 |
| Cell Dimension | |
| a, b, c (Å) | 110.09, 52.10, 44.47 |
| α, β, γ (°) | 90.00, 103.29, 90.00 |
| Resolution (Å) | 50.00 – 2.60 |
| | (2.64 – 2.60) |
| $R_{merge}$ | 0.109 (0.613) |
| <I>/σ<I> | 11.76 (2) |
| Completeness (%) | 97.7 (98.5) |
| Redundancy | 3.5 (3.4) |
| **Refinement** | |
| Resolution (Å) | 50 – 2.60 |
| | (2.664 - 2.60) |
| No. reflections/free | 7460 / 779 |
| $R_{work}/R_{free}$ | 0.210/ 0.279 |
| No. Atoms | 2398 |
| Protein | 2340 |
| Ligand/Ion | 50 |
| Water | 8 |
| B-Factors ($Å^2$) | |
| Protein | 59.25 |
| Ligand/Ion | 61.04 |
| Solvent | 49.14 |
| RMS Deviations | |
| Bond Lengths (Å) | 0.013 |
| Bond Angles (°) | 1.76 |
| Ramachandran Favored (%) | 97.01 |
| Ramachandran Allowed (%) | 2.99 |
| Ramachandran Outliers (%) | 0.00 |
| Rotameric Outliers (%) | 1.92 |
| Clashscore | 11.06 |

### Compounds synthesis and characterization

General chemical methods. All chemicals were purchased from commercial vendors and used without further purification unless otherwise noted. $^1$H and $^{13}$C NMR spectra were recorded on a Bruker-400 or -500 NMR spectrometer. Chemical shifts are reported in parts per million referenced with respect to residual solvent ($CD_3OD$) 3.31 ppm, (DMSO-d6) 2.50 ppm, and ($CDCl_3$) 7.26 ppm or from internal standard tetramethylsilane (TMS) 0.00 ppm. The following abbreviations were used in reporting spectra: s, singlet; d, doublet; t, triplet; q, quartet; m, multiplet; dd, doublet of doublets; ddd, doublet of doublet of doublets. All reactions were carried out under $N_2$ atmosphere, unless otherwise stated. HPLC-grade solvents were used for all reactions. Flash column chromatography was performed using silica gel (230–400 mesh, Merck). Low-resolution mass spectra were obtained using an ESI technique on a 3200 Q Trap LC/MS/MS system (Applied Biosystems). The purity was assessed by using Shimadzu LC-MS with Waters XTerra MS C-18 column (part #186000538), 50 × 2.1 mm, at a flow rate of 0.3 mL/min; λ = 250 and 220 nm; mobile phase A, 0.1% formic acid in $H_2O$, and mobile phase B', 0.1% formic in 60% isopropanol, 30% $CH_3CN$ and 9.9% $H_2O$. All compounds submitted for testing in TEVC assay and plaque reduction assay were confirmed to be > 95.0% purity by LC-MS traces. All final products were characterized by proton and carbon NMR and MS.

General Procedure for Ugi-4CR Reaction

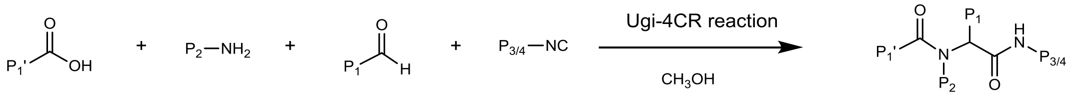

Amine (1.0 equiv) and aldehyde (1.0 equiv) were mixed in methanol (10 ml) and stirred at room temperature for 30 minutes. Then carboxylic acid (1.0 equiv) and isocyanide (1.0 equiv) were added sequentially and the resulting mixture was stirred at room temperature overnight. After that, the solvent was removed under reduced pressure and the crude product was purified with flash silica gel chromatography (methanol in dichloromethane 1-5% or acetone in hexane 30-80%).

<u>General procedures for the synthesis of compounds **24, 28, 33, 35, and 37** by TFA deprotection</u>

To a solution of N-Trityl-protected Ugi-4CR compound in dichloromethane (5 mL) was added TFA (1 mL). The mixture was stirred at room temperature for 2 h, and the solvent was removed under reduced pressure. The crude mixture was diluted in dichloromethane and purified by silica gel flash column chromatography (ammonia 10 % methanol in dichloromethane 10 – 15 %) to give the final product.

<u>Procedure for the synthesis of compound **18** by Suzuki -Miyamura Cross-coupling</u>

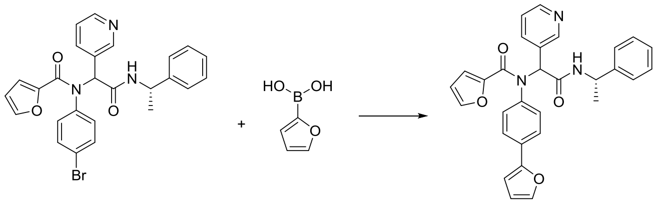

**18**

To solution of 2-[N-(4-bromophenyl)-1-(furan-2-yl)formamido]-N-[(1S)-1-phenylethyl]-2-(pyridin-3-yl)acetamide (1 mmol) and furan-2-boronic acid (1 mmol) in 1,4-dioxane in a microwave reaction vial was added an aqueous solution of $K_2CO_3$ (4 mmol). The resulting solution was purged with $N_2$ for 5 min. The catalyst, $Pd(PPh_3)_4$ (0.1 mmol), was added in one portion. The vial was capped and heated to 140 $^oC$ for 30 minutes with microwave irradiation. After cooling down to room temperature, the reaction solution was diluted with dichloromethane and extracted with water, followed by brine. The organic layer was dried over $MgSO_4$, filtrated, and concentrated under reduced pressure. The crude product was purified by silica gel flash column chromatography (Methanol in dichloromethane 1-5%) to give the final product.

Procedure for the synthesis of 4-(thiophen-2-yl)aniline by Suzuki -Miyamura Cross-coupling

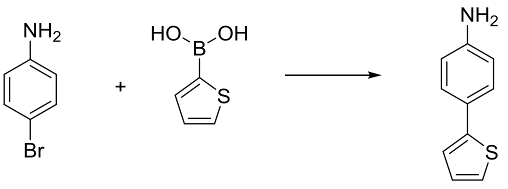

The starting material 4-(thiophen-2-yl)aniline used for the synthesis of compound **19** was prepared using the following procedure. To suspension of thiophene-2-boronic acid (1.0 mmol), sodium carbonate (1.0 mmol) in toluene/methanol (4:1, 40 mL) was added 4-bromoaniline (1.0 mmol). The resulting solution was purged with $N_2$ for 10 min. The catalyst, $Pd(PPh_3)_4$ (0.1 mmol), was added in one portion. The reaction mixture was stirred for 16 hr. at 100 °C. After cooling to room temperature, the reaction mixture was diluted with ethyl acetate (100 mL) and washed with water (100 mL). The organic layer was dried over $MgSO_4$, filtrated, and concentrated under reduced pressure. The crude product was purified by silica gel flash column chromatography (Ethyl acetate in Hexane 20 – 40 %) to give the final product.

4-(thiophen-2-yl)aniline. $^1H$ NMR (500 MHz, $CDCl_3$) δ 7.43 – 7.38 (m, 2H), 7.17 – 7.12 (m, 2H), 7.01 (dd, $J$ = 5.0, 3.6 Hz, 1H), 6.69 – 6.63 (m, 2H), 3.68 (s, 2H). $^{13}C$ NMR (126 MHz, $CDCl_3$) δ 146.10, 145.11, 127.93, 127.25, 125.24, 123.17, 121.36, 115.39. Yield. 31.2 %.

The isocyanide used for the synthesis of compound **40** was prepared used the method below:

Procedure for synthesis of N-(3-phenylpropyl)formamide

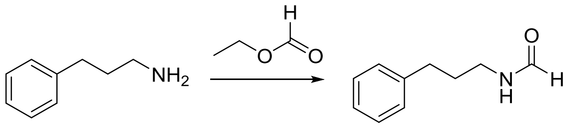

3-phenyl propylamine (2 mmol) in a microwave reaction vial was mixed with ethyl formate (6 mL). The vial was capped and heated to 70 °C overnight. After reaction residual ethyl formate was evaporated under reduced pressure. The crude product was purified by silica gel flash column chromatography (methanol in dichloromethane 1 - 3 %) to give the product N-(3-phenylpropyl)formamide. $^{1}$H NMR (500 MHz, $CDCl_3$) δ 8.15 – 7.96 (m, 1H), 7.33 – 7.24 (m, 2H), 7.24 – 7.12 (m, 3H), 6.18 – 5.73 (m, 1H), 3.35 – 3.17 (m, 2H), 2.69 – 2.60 (m, 2H), 1.89 – 1.78 (m, 2H). $^{13}$C NMR (126 MHz, $CDCl_3$) δ 170.33, 164.88, 161.44, 141.51, 141.27, 140.65, 128.67, 128.55, 128.52, 128.42, 128.41, 126.32, 126.13, 126.07, 41.15, 39.39, 37.84, 33.36, 33.22, 32.60, 32.54, 31.17, 31.16, 23.31. Yield. 98.1%.

Procedure for the synthesis of compound **40**

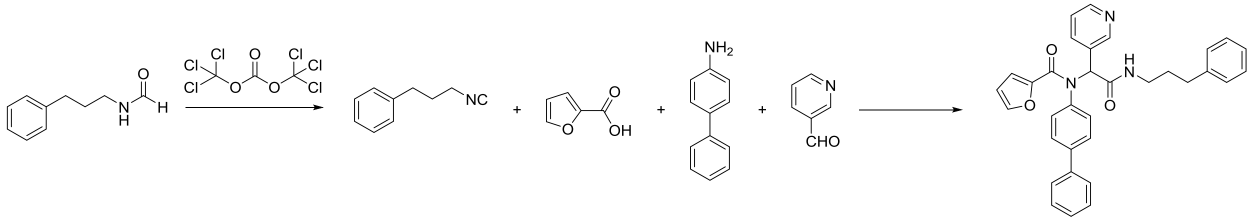

40

To solution of N-(3-phenylpropyl)formamide (2 mmol) in dichloromethane (50 mL) was added triethylamine (4.8 mmol). The solution was cooled to approximately – 10 °C using an ethanol ice bath. Triphosgene (0.4 mmol) was added to the stirring mixture. The resulting mixture was stirred at – 10 °C for 10 minutes. In separate 250 ml round-bottomed flask, 4-aminobiphneyl (2 mmol) and 3-pyridinecarboxaldehyde (2 mmol) were mixed in methanol (100 ml) and stirred at room temperature for 30 minutes. Then carboxylic acid (2 mmol) and the resulting isocyanide solution (1.0 equiv) were added sequentially and the resulting mixture was stirred at room temperature overnight. After that, the solvent was removed under reduced pressure and the crude product was purified with flash silica gel chromatography (Methanol in dichloromethane 1-5%) to give the final product.

Characterizations of the final products:

N-tert-butyl-2-[N-(4-tert-butylphenyl)-1-(furan-2-yl)formamido]-2-(pyridin-3-yl)acetamide (**1**). $^1$H NMR (400 MHz, DMSO-$d_6$) δ 8.38 – 8.24 (m, 2H), 7.86 (s, 1H), 7.69 – 7.66 (m, 1H), 7.37 (dt, $J$ = 7.9, 2.0 Hz, 1H), 7.26 – 7.17 (m, 2H), 7.26 – 7.02 (br s, 1H), 7.14 – 7.04 (m, 2H), 6.31 (dd, $J$ = 3.6, 1.7 Hz, 1H), 6.15 (s, 1H), 5.24 (d, $J$ = 3.5 Hz, 1H), 1.22 (s, 9H), 1.19 (s, 9H). $^{13}$C NMR (101 MHz, DMSO-$d_6$) δ 168.07, 158.14, 151.12, 150.95, 148.53, 146.37, 145.18, 137.26, 136.53, 131.23, 130.83, 125.20, 122.73, 115.75, 111.24, 62.30, 50.41, 34.27, 31.01, 28.32. Yield: 81.2%. $C_{26}H_{31}N_3O_3$, HRMS calculated for m/z $[M+H]^+$: 434.244367, found 434.2438.

2-[N-(4-tert-butylphenyl)-1-(furan-2-yl)formamido]-N-(propan-2-yl)-2-(pyridin-3-yl)acetamide (**2**). $^1$H NMR (400 MHz, DMSO-d$_6$) δ 8.37 – 8.28 (m, 2H), 8.10 (d, $J$ = 7.5 Hz, 1H), 7.76 – 7.61 (m, 1H), 7.37 (dt, $J$ = 7.9, 2.0 Hz, 1H), 7.30 – 7.17 (m, 2H), 7.24 – 7.00 (br s, 1H), 7.17 – 7.07 (m, 2H), 6.30 (dd, $J$ = 3.6, 1.7 Hz, 1H), 6.15 (s, 1H), 5.26 (d, $J$ = 3.6 Hz, 1H), 3.99 – 3.76 (m, 1H), 1.20 (s, 9H), 1.07 (d, $J$ = 6.6 Hz, 3H), 0.95 (d, $J$ = 6.5 Hz, 3H). $^{13}$C NMR (101 MHz, DMSO-d$_6$) δ 167.69, 158.23, 151.15, 151.05, 148.69, 146.36, 145.27, 137.41, 136.52, 131.01, 130.77, 125.28, 122.81, 115.85, 111.27, 62.10, 40.85, 34.30, 31.02, 22.09, 22.07.

Yield. 82.3%. $C_{25}H_{29}N_3O_3$, HRMS calculated for m/z $[M+H]^+$: 420.228716 (calculated), 420.2282 (found).

2-[N-(4-tert-butylphenyl)-1-(furan-2-yl)formamido]-N-cyclopropyl-2-(pyridin-3-yl)acetamide (**3**). $^1H$ NMR (400 MHz, DMSO-$d_6$) δ 8.34 – 8.28 (m, 3H), 7.67 (dd, *J* = 1.7, 0.7 Hz, 1H), 7.37 (dt, *J* = 8.0, 1.9 Hz, 1H), 7.26 – 7.19 (m, 2H), 7.25 – 6.94 (br s, 1H ), 7.16 – 7.09 (m, 2H), 6.31 (dd, *J* = 3.6, 1.7 Hz, 1H), 6.09 (s, 1H), 5.28 (dd, *J* = 3.6, 0.8 Hz, 1H), 2.71 – 2.60 (m, 1H), 1.20 (s, 9H), 0.67 – 0.53 (m, 2H), 0.40 – 0.25 (m, 2H). $^{13}C$ NMR (101 MHz, DMSO-$d_6$) δ 169.86, 158.25, 151.10, 148.75, 146.31, 145.29, 137.40, 136.46, 130.70, 125.31, 122.84, 115.44, 111.27, 62.05, 34.30, 31.01, 22.02, 5.17. Yield. 80.8%. $C_{25}H_{27}N_3O_3$, HRMS calculated for m/z $[M+H]^+$: 418.213067 (calculated), 418.2125 (found).

2-[N-(4-tert-butylphenyl)-1-(furan-2-yl)formamido]-2-(pyridin-3-yl)-N-(2,4,4-trimethylpentan-2-yl)acetamide (**4**). $^1H$ NMR (500 MHz, DMSO-$d_6$) δ 8.36 – 8.28 (m, 2H), 7.79 – 7.59 (m, 2H), 7.43 – 7.35 (m, 1H), 7.32-6.97 (br s, 1H), 7.30 – 7.18 (m, 2H), 7.12 (dd, *J* = 8.0, 4.8 Hz, 1H), 6.36 – 6.26 (m, 1H), 6.18 (s, 1H), 5.25 (d, *J* = 3.6 Hz, 1H), 1.67 (dd, *J* = 75.5, 14.6 Hz, 2H), 1.35 – 1.23 (m, 7H), 1.21 (s, 9H), 0.86 (s, 9H). $^{13}C$ NMR (101 MHz, DMSO-$d_6$) δ 167.63, 158.09, 151.19, 150.99, 148.52, 146.39, 145.14, 137.41, 136.62, 131.04, 130.88, 125.22, 122.58, 115.72, 111.22, 62.45, 54.36, 50.65, 34.27, 31.16, 31.12, 31.10, 31.01, 28.99, 28.37.
Yield. 60.0%. $C_{30}H_{39}N_3O_3$, HRMS calculated for m/z $[M+H]^+$: 490.306967 (calculated), 490.3064 (found).

2-[N-(4-tert-butylphenyl)-1-(furan-2-yl)formamido]-N-cyclopentyl-2-(pyridin-3-yl)acetamide (**5**). $^1H$ NMR (400 MHz, DMSO-$d_6$) δ 8.36 – 8.28 (m, 2H), 8.18 (d, *J* = 7.0 Hz, 1H), 7.70 – 7.65 (m, 1H), 7.37 (dt, *J* = 7.9, 2.0 Hz, 1H), 7.28 – 7.18 (m, 2H), 7.25 – 6.94 (br s, 1H) 7.18 – 7.06

(m, 2H), 6.30 (dd, *J* = 3.6, 1.7 Hz, 1H), 6.16 (s, 1H), 5.26 (d, *J* = 3.6 Hz, 1H), 4.02 (h, *J* = 6.9 Hz, 1H), 1.86 – 1.67 (m, 2H), 1.65 – 1.31 (m, 5H), 1.30 – 1.21 (m, 1H), 1.20 (s, 9H). $^{13}$C NMR (101 MHz, DMSO-$d_6$) δ 168.13, 158.24, 151.12, 151.04, 148.67, 146.36, 145.26, 137.35, 136.51, 131.02, 130.77, 125.27, 122.80, 115.85, 111.27, 62.04, 50.69, 34.30, 32.05, 31.86, 31.02, 23.48, 23.43. Yield. 85.8%. $C_{27}H_{31}N_3O_3$, HRMS calculated for m/z $[M+H]^+$: 446.244367 (calculated), 446.2438 (found).

2-[N-(4-tert-butylphenyl)-1-(furan-2-yl)formamido]-N-cyclohexyl-2-(pyridin-3-yl)acetamide (**6**). $^1$H NMR (500 MHz, DMSO-$d_6$) δ 8.36 – 8.30 (m, 2H), 8.09 (d, *J* = 7.7 Hz, 1H), 7.67 (dd, *J* = 1.7, 0.7 Hz, 1H), 7.38 (dt, *J* = 7.9, 2.0 Hz, 1H), 7.28 – 7.02 (br s, 1H), 7.25 – 7.19 (m, 2H), 7.17 – 7.10 (m, 2H), 6.31 (dd, *J* = 3.6, 1.7 Hz, 1H), 6.18 (s, 1H), 5.28 (d, *J* = 3.6 Hz, 1H), 3.63 – 3.52 (m, 1H), 1.86 – 1.40 (m, 5H), 1.32 – 1.14 (s, 12H), 1.14 – 0.92 (m, 2H). $^{13}$C NMR (101 MHz, DMSO-$d_6$) δ 167.66, 157.96, 151.12, 151.01, 148.63, 146.37, 145.23, 137.35, 136.53, 131.08, 130.75, 125.24, 122.78, 116.21, 111.24, 62.07, 47.98, 34.28, 32.07, 31.00, 25.15, 24.51, 24.39. Yield. 81.7%. $C_{28}H_{33}N_3O_3$, HRMS calculated for m/z $[M+H]^+$: 460.260017 (calculated), 460.2595 (found).

2-[N-(4-tert-butylphenyl)-1-(furan-2-yl)formamido]-N-(4-fluorophenyl)-2-(pyridin-3-yl)acetamide (**7**). $^1$H NMR (400 MHz, MeOD-$d_4$) δ 8.43 (d, *J* = 2.3 Hz, 1H), 8.34 (dd, *J* = 4.9, 1.6 Hz, 1H), 7.63 (dt, *J* = 8.0, 2.0 Hz, 1H), 7.59 – 7.53 (m, 2H), 7.52 (d, *J* = 1.8 Hz, 1H), 7.45 – 6.60 (br s, 2H), 7.30 (s, 2H), 7.25 – 7.17 (m, 1H), 7.08 – 6.98 (m, 2H), 6.37 (s, 1H), 6.25 (dd, *J* = 3.6, 1.7 Hz, 1H), 5.46 (d, *J* = 3.6 Hz, 1H), 1.26 (s, 9H). $^{13}$C NMR (101 MHz, MeOD-$d_4$) δ 169.50, 161.43, 153.85, 152.20, 149.96, 147.43, 146.74, 140.32, 139.46, 137.47, 134.59, 132.24,

131.96, 127.10, 125.58, 123.10, 123.02, 118.43, 116.44, 116.22, 112.35, 65.57, 35.53, 31.62. Yield. 86.1 %. $C_{28}H_{26}FN_3O_3$, HRMS calculated for m/z $[M+H]^+$: 472.203645 (calculated), 472.2031 (found).

2-[N-(4-tert-butylphenyl)-1-(furan-2-yl)formamido]-N-(4-methoxyphenyl)-2-(pyridin-3-yl)acetamide (**8**). $^1$H NMR (400 MHz, DMSO-$d_6$) δ 10.22 (s, 1H), 8.42 (d, *J* = 2.3 Hz, 1H), 8.34 (dd, *J* = 4.8, 1.6 Hz, 1H), 7.72 – 7.67 (m, 1H), 7.54 – 7.46 (m, 2H), 7.43 (dt, *J* = 8.0, 2.0 Hz, 1H), 7.35 – 7.05 (br s, 2H), 7.27 – 7.21 (m, 2H), 7.17 – 7.10 (m, 1H), 6.92 – 6.83 (m, 2H), 6.32 (dd, *J* = 3.6, 1.7 Hz, 1H), 6.30 (s, 1H), 5.29 (d, *J* = 3.6 Hz, 1H), 3.71 (s, 3H), 1.20 (s, 9H). $^{13}$C NMR (101 MHz, DMSO-$d_6$) δ 167.33, 158.37, 155.38, 151.34, 151.25, 149.02, 146.23, 145.45, 137.54, 136.32, 131.99, 130.87, 130.21, 125.42, 122.95, 120.71, 116.06, 113.91, 111.82, 62.78, 55.18, 34.33, 31.02. Yield. 79.1 %. $C_{29}H_{29}N_3O_4$, HRMS calculated for m/z $[M+H]^+$: 484.223632 (calculated), 484.2231 (found).

2-[N-(4-tert-butylphenyl)-1-(furan-2-yl)formamido]-N-(2,6-dimethylphenyl)-2-(pyridin-3-yl)acetamide (**9**). $^1$H NMR (400 MHz, DMSO-$d_6$) δ 9.66 (s, 1H), 8.48 (s, 1H), 8.37 (d, *J* = 4.9 Hz, 1H), 7.70 (dd, *J* = 1.7, 0.7 Hz, 1H), 7.50 (dt, *J* = 8.0, 2.0 Hz, 1H), 7.28 – 7.08 (br s, 2H), 7.28 – 7.21 (m, 2H), 7.21 – 7.13 (m, 1H), 7.10 – 6.99 (m, 3H), 6.35 (s, 1H), 6.32 (dd, *J* = 3.6, 1.7 Hz, 1H), 5.28 (dd, *J* = 3.6, 0.8 Hz, 1H), 2.07 (s, 6H), 1.21 (s, 9H). $^{13}$C NMR (101 MHz, DMSO-$d_6$) δ δ 167.74, 158.30, 151.46, 151.24, 149.09, 146.28, 145.31, 137.71, 136.44, 135.44, 134.60, 130.87, 130.22, 127.60, 126.53, 125.42, 122.75, 115.90, 111.29, 62.64, 34.31, 31.00, 17.99. Yield. 75.9 %. $C_{30}H_{31}N_3O_3$, HRMS calculated for m/z $[M+H]^+$: 482.244367 (calculated), 482.2438 (found).

N-[(1H-1,2,3-benzotriazol-1-yl)methyl]-2-[N-(4-tert-butylphenyl)-1-(furan-2-yl)formamido]-2-(pyridin-3-yl)acetamide (**10**). $^{1}$H NMR (500 MHz, $CDCl_3$) δ 8.80 (t, *J* = 6.7 Hz, 1H), 8.50 – 8.46 (m, 1H), 8.40 – 8.36 (m, 1H), 8.10 (d, *J* = 8.3 Hz, 1H), 8.02 (d, *J* = 8.6 Hz, 1H), 7.63 – 7.55 (m, 1H), 7.52 – 7.38 (m, 3H), 7.24 (s, 1H), 7.12 – 6.83 (m, 3H), 6.38 (s, 1H), 6.33 – 6.20 (m, 3H), 5.47 – 5.40 (m, 2H), 1.37 (s, 9H). $^{13}$C NMR (126 MHz, $CDCl_3$) δ 169.85, 159.85, 152.70, 151.60, 149.74, 146.04, 145.97, 145.30, 138.30, 136.13, 132.52, 130.17, 129.66, 127.93, 126.20, 124.40, 122.96, 119.63, 117.55, 111.38, 111.08, 63.02, 51.48, 34.78, 31.34.
Yield. 22.3 %. $C_{29}H_{28}N_6O_3$, HRMS calculated for m/z $[M+H]^+$: 509.230114 (calculated), 509.2296 (found).

2-[N-(4-tert-butylphenyl)-1-(furan-2-yl)formamido]-N-[(4-methylbenzenesulfonyl)methyl]-2-(pyridin-3-yl)acetamide (**11**). $^{1}$H NMR (400 MHz, DMSO-$d_6$) δ 9.28 – 9.20 (m, 1H), 8.37 (dd, *J* = 4.3, 2.2 Hz, 1H), 8.23 (s, 1H), 7.69 (dd, *J* = 1.8, 0.7 Hz, 1H), 7.56 – 7.49 (m, 2H), 7.30 – 7.26 (m, 2H), 7.24 – 7.17 (m, 2H), 7.15 – 7.05 (m, 2H), 6.96 (s, 2H), 6.31 (dd, *J* = 3.6, 1.7 Hz, 1H), 6.23 (s, 1H), 5.26 (d, *J* = 3.6 Hz, 1H), 4.93 (dd, *J* = 14.1, 7.5 Hz, 1H), 4.61 (dd, *J* = 14.1, 5.6 Hz, 1H), 2.34 (s, 3H), 1.20 (s, 9H). $^{13}$C NMR (101 MHz, DMSO-$d_6$) δ 169.31, 158.16, 151.48, 151.25, 148.92, 146.15, 145.42, 144.46, 137.74, 136.09, 134.47, 130.70, 129.59, 128.43, 125.32, 122.62, 116.01, 111.81, 61.64, 60.19, 34.30, 30.99, 21.06. Yield. 49.9 %. $C_{30}H_{31}N_3O_5S$, HRMS calculated for m/z $[M+H]^+$: 546.206269 (calculated), 546.2057 (found).

N-benzyl-2-[N-(4-tert-butylphenyl)-1-(furan-2-yl)formamido]-2-(pyridin-3-yl)acetamide (**12**). $^{1}$H NMR (400 MHz, DMSO-$d_6$) δ 8.74 (t, *J* = 5.9 Hz, 1H), 8.37 – 8.31 (m, 2H), 7.72 – 7.66 (m, 1H), 7.39 (dt, *J* = 8.0, 1.9 Hz, 1H), 7.33 – 7.18 (m, 7H), 7.21 – 7.05 (br s, 1H), 7.17 – 7.08 (m,

2H), 6.31 (dd, $J$ = 3.6, 1.7 Hz, 1H), 6.22 (s, 1H), 5.28 (d, J =4.21 Hz, 1H), 4.42 – 4.27 (m, 2H), 1.21 (s, 9H). $^{13}$C NMR (101 MHz, DMSO-d$_6$) $^{13}$C NMR (101 MHz, DMSO) δ 168.92, 158.28, 151.32, 151.15, 148.84, 146.31, 145.30, 139.19, 137.63, 136.57, 130.70, 128.18, 127.14, 126.74, 125.38, 122.74, 115.93, 111.28, 62.48, 42.34, 34.30, 31.00. Yield. 49.9 %. $C_{29}H_{29}N_3O_3$, HRMS calculated for m/z $[M+H]^+$: 468.228716 (calculated), 468.2282 (found).

2-[N-(4-tert-butylphenyl)-1-(furan-2-yl)formamido]-N-[(1S)-1-phenylethyl]-2-(pyridin-3-yl)acetamide (**13**). $^1$H NMR (400 MHz, DMSO-D$_6$) δ 8.68 (dd, $J$ = 7.8, 2.7 Hz, 1H), 8.41 – 8.23 (m, 2H), 7.67 (ddd, $J$ = 5.7, 1.7, 0.7 Hz, 1H), 7.46 – 7.24 (m, 3H), 7.28 – 6.94 (br s, 1H), 7.23 – 7.11 (m, 5H), 7.11 – 7.01 (m, 2H), 6.34 – 6.24 (m, 2H), 5.28 (dd, $J$ = 3.6, 0.8 Hz, 0.5H), 5.22 (dd, $J$ = 3.6, 0.8 Hz, 0.5H), 5.04 – 4.91 (m, 1H), 1.37 (d, $J$ = 7.0 Hz, 1.5H), 1.25 (d, $J$ = 7.0 Hz, 1.5H), 1.19 (d, $J$ = 6.0 Hz, 9H). $^{13}$C NMR (101 MHz, DMSO) δ 168.14, 167.92, 158.27, 158.18, 151.31, 151.26, 151.06, 148.76, 146.35, 146.28, 145.28, 145.24, 144.30, 144.00, 137.51, 137.49, 136.41, 136.39, 130.80, 130.46, 128.19, 128.06, 126.65, 126.58, 126.16, 125.75, 125.26, 122.83, 122.53, 115.87, 115.83, 111.25, 111.24, 62.01, 48.34, 48.20, 34.26, 31.00, 30.98, 22.26, 22.10. Yield. 84.2 %. $C_{30}H_{31}N_3O_3$, HRMS calculated for m/z $[M+H]^+$: 482.244367 (calculated), 482.2438 (found).

2-[1-(furan-2-yl)-N-[4-(trimethylsilyl)phenyl]formamido]-N-[(1S)-1-phenylethyl]-2-(pyridinyl)acetamide (**14**). $^1$H NMR (500 MHz, $CDCl_3$) δ 8.50 – 8.41 (m, 2H), 7.47 (dd, $J$ = 49.6, 8.2 Hz, 1H), 7.39 – 7.16 (m, 8H), 7.10 – 6.81 (m, 4H), 6.25 (d, $J$ = 13.4 Hz, 1H), 6.18 – 6.13 (m, 1H), 5.50 – 5.45 (m, 0.5H), 5.44 – 5.39 (m, 0.5H), 5.22 – 5.12 (m, 1H), 1.54 (d, $J$ = 6.9 Hz, 1.5H), 1.48 (d, $J$ = 7.0 Hz, 1.5H), 0.23 (s, 9H). $^{13}$C NMR (126 MHz, $CDCl_3$) δ 168.01, 167.79, 159.83, 159.76, 151.55, 151.47, 149.82, 149.75, 146.29, 146.21, 145.19, 145.14, 143.14,

142.17, 142.06, 139.73, 139.56, 138.42, 138.16, 134.17, 134.16, 130.21, 130.15, 130.01, 129.86, 128.77, 128.68, 127.42, 127.34, 126.34, 126.14, 122.98, 122.96, 117.46, 117.44, 111.37, 111.35, 63.46, 62.97, 49.54, 49.51, 22.06, 21.92, 0.10. Yield. 50.2 %. $C_{29}H_{31}N_3O_3Si$, HRMS calculated for m/z $[M+H]^+$: 498.221295 (calculated), 498.2220 (found).

2-{N-[4-(tert-butoxy)phenyl]-1-(furan-2-yl)formamido}-N-[(1S)-1-phenylethyl]-2-(pyridin-3-yl)acetamide (**15**). $^1$H NMR (500 MHz, DMSO-$d_6$) δ 8.72 (t, *J* = 8.17 Hz, 1H), 8.31 (dd, J = 55, 2.4 Hz, 1H), 8.33 (ddd, *J* = 13.5, 4.8, 1.6 Hz, 1H), 7.69 (ddd, *J* = 7.0, 1.7, 0.8 Hz, 1H), 7.50-7.00 (br s, 1H), 7.42-7.06 (m, 3H), 7.43 – 7.32 (m, 1H), 7.31 – 7.12 (m, 3H), 6.76 (s, 3H), 6.34 (ddd, *J* = 8.2, 3.6, 1.7 Hz, 1H), 6.27 (d, *J* = 13.2 Hz, 1H), 5.46 (d, *J* = 3.6 Hz, 0.5H), 5.41 (d, *J* = 3.6 Hz, 0.5H), 5.05 – 4.94 (m, 1H), 1.38 (d, *J* = 7.1 Hz, 1.5H), 1.25 (d, *J* = 7.0 Hz, 1.5H), 1.19 (d, *J* = 9.5 Hz, 9H). $^{13}$C NMR (126 MHz, DMSO-$d_6$) δ 168.32, 168.10, 166.39, 158.22, 158.14, 154.78, 154.76, 151.49, 151.44, 148.78, 146.45, 146.39, 145.35, 145.30, 144.36, 144.03, 137.68, 137.65, 134.18, 134.14, 132.04, 130.75, 130.38, 128.26, 128.13, 126.72, 126.65, 126.20, 125.77, 123.73, 122.91, 122.60, 116.11, 116.07, 111.26, 111.24, 78.39, 62.08, 48.40, 48.26, 28.43, 22.38, 22.16. Yield. 75.2 %. $C_{30}H_{31}N_3O_4$, HRMS calculated for m/z $[M+H]^+$: 498.239282 (calculated), 498.2387 (found).

2-[1-(furan-2-yl)-N-[4-(piperidin-1-yl)phenyl]formamido]-N-[(1S)-1-phenylethyl]-2-(pyridin-3-yl)acetamide (**16**). $^1$H NMR (400 MHz, DMSO-$d_6$) δ 8.65 (dd, *J* = 7.8, 3.6 Hz, 1H), 8.40 – 8.19 (m, 2H), 7.70 (ddd, *J* = 6.3, 1.7, 0.7 Hz, 1H), 7.64-7.01 (br s, 1H), 7.48 – 7.29 (m, 3H), 7.28 – 7.03 (m, 4H), 6.68 (s, 3H), 6.35 – 6.28 (m, 1H), 6.25 (d, *J* = 7.8 Hz, 1H), 5.30 – 5.23 (m, 0.5H), 5.22 – 5.13 (m, 0.5H), 4.98 (h, *J* = 7.1 Hz, 1H), 3.11 – 3.02 (m, 4H), 1.53 (s, 6H), 1.37 (d, *J* = 7.0 Hz, 1.5H), 1.24 (d, *J* = 7.0 Hz, 1.5H). $^{13}$C NMR (101 MHz, DMSO-$d_6$) δ 168.34, 168.11,

158.58, 158.51, 151.36, 151.29, 150.81, 148.76, 146.41, 146.36, 145.19, 145.14, 144.36, 144.05, 137.67, 137.63, 131.74, 128.80, 128.77, 128.19, 128.07, 126.64, 126.58, 126.17, 125.75, 115.78, 115.74, 114.55, 111.30, 62.08, 54.90, 48.68, 48.31, 48.18, 30.68, 25.76, 24.94, 24.91, 24.16, 23.78, 22.32. Yield. 78.2 %. $C_{31}H_{32}N_4O_3$, HRMS calculated for m/z $[M+H]^+$: 509.255266 (calculated), 509.2547 (found).

2-[N-(4-cyclohexylphenyl)-1-(furan-2-yl)formamido]-N-[(1S)-1-phenylethyl]-2-(pyridin-3-yl)acetamide (**17**). $^1$H NMR (400 MHz, DMSO-$d_6$) δ 8.68 (dd, *J* = 7.7, 3.4 Hz, 1H), 8.32 (dd, *J* = 10.5, 4.7 Hz, 1H), 8.31 (d, J = 48 Hz, 1H), 7.70 – 7.64 (m, 1H), 7.44 – 7.28 (m, 3H), 7.28 – 7.11 (m, 4H), 7.10 – 6.98 (m, 4H),6.33 – 6.24 (m, 2H), 5.25 (d, *J* = 3.6 Hz, 0.5H), 5.20 (d, *J* = 3.6 Hz, 0.5H), 5.05 – 4.91 (m, 1H), 2.48 – 2.31 (m, 1H), 1.84 – 1.57 (m, 5H), 1.39-1.10 (m, 5H), 1.31 (dd, J = 41.17, 7.02 Hz, 3H) $^{13}$C NMR (101 MHz, DMSO-$d_6$) δ 168.18, 167.96, 158.25, 158.17, 151.34, 151.29, 148.78, 147.88, 146.34, 146.28, 145.28, 145.24, 144.30, 144.00, 137.51, 136.68, 136.66, 131.12, 128.19, 128.07, 126.73, 126.66, 126.59, 126.16, 125.75, 115.88, 115.84, 111.24, 62.03, 48.34, 48.21, 43.10, 33.85, 33.81, 33.77, 26.18, 25.47, 22.27, 22.10. Yield. 83.4 %. $C_{32}H_{33}N_3O_3$, HRMS calculated for m/z $[M+H]^+$: 508.260017 (calculated), 508.2595 (found).

2-[1-(furan-2-yl)-N-[4-(furan-2-yl)phenyl]formamido]-N-[(1S)-1-phenylethyl]-2-(pyridin-3-yl)acetamide (**18**). $^1$H NMR (500 MHz, DMSO-$d_6$) δ 8.80 (dd, *J* = 13.9, 7.8 Hz, 1H), 8.66 – 8.45 (m, 2H), 7.82 (dt, *J* = 8.1, 1.9 Hz, 1H), 7.71 (dddd, *J* = 24.9, 6.9, 1.8, 0.8 Hz, 2H), 7.64 – 7.46 (m, 3H), 7.45 – 7.07 (m, 7H), 6.96 (ddd, *J* = 5.9, 3.4, 0.8 Hz, 1H), 6.62 – 6.55 (m, 1H), 6.38 (d, *J* = 11.7 Hz, 1H), 6.35 (ddd, *J* = 8.2, 3.6, 1.7 Hz, 1H), 5.66 (d, *J* = 2.8 Hz, 0.5H), 5.62 (d, *J* = 3.6, 0.8 Hz, 0.5H), 5.06 – 4.94 (m, 1H), 1.37 (d, *J* = 7.0 Hz, 1.5H), 1.29 (d, *J* = 7.0 Hz, 1.5H). $^{13}$C NMR (126 MHz, DMSO-$d_6$) δ 167.42, 167.26, 158.28, 158.22, 151.98, 148.37, 148.26, 146.21,

146.16, 145.56, 145.52, 144.16, 143.82, 143.41, 141.34, 141.18, 138.11, 138.07, 132.42, 132.05, 131.68, 131.64, 130.08, 128.24, 128.16, 126.75, 126.69, 126.19, 125.82, 124.47, 124.16, 123.58, 116.64, 116.61, 112.26, 111.45, 111.44, 107.04, 61.95, 48.52, 48.37, 22.22, 21.99. Yield. 35.2 %. $C_{30}H_{25}N_3O_4$, HRMS calculated for m/z $[M+H]^+$: 492.192331 (calculated), 492.1918 (found).

2-[1-(furan-2-yl)-N-[4-(thiophen-2-yl)phenyl]formamido]-N-[(1S)-1-phenylethyl]-2-(pyridin-3-yl)acetamide (**19**). $^1$H NMR (500 MHz, DMSO-$d_6$) δ 8.79 – 8.71 (m, 1H), 8.48 – 8.23 (m, 2H), 7.74 – 7.64 (m, 1H), 7.58 – 7.45 (m, 4H), 7.44 – 7.31 (m, 3H), 7.30 – 7.14 (m, 4H), 7.14 – 7.05 (m, 3H), 6.40 – 6.28 (m, 2H), 5.63 – 5.69 (m, 0.5H), 5.58 – 5.54 (m, 0.5H), 5.07 – 4.96 (m, 1H), 1.39 (d, *J* = 7.1 Hz, 1.5H), 1.26 (d, *J* = 7.0 Hz, 1.5H). $^{13}$C NMR (126 MHz, DMSO-$d_6$) δ 168.15, 167.96, 158.21, 158.14, 151.40, 151.33, 148.97, 146.31, 146.25, 145.42, 145.37, 144.27, 143.98, 142.06, 138.29, 138.25, 137.67, 137.63, 133.43, 133.34, 133.33, 131.98, 130.66, 130.28, 128.60, 128.58, 128.22, 128.10, 126.68, 126.63, 126.29, 126.16, 126.14, 125.78, 125.15, 124.47, 124.45, 123.09, 122.80, 116.36, 116.33, 111.39, 111.37, 62.02, 62.00, 48.39, 48.25, 22.30, 22.13. Yield. 81.1%. $C_{30}H_{25}N_3O_3S$, HRMS calculated for m/z $[M+H]^+$: 508.169489 (calculated), 508.1689 (found).

2-[1-(furan-2-yl)-N-[4-(1H-pyrrol-1-yl)phenyl]formamido]-N-[(1S)-1-phenylethyl]-2-(pyridin-3-yl)acetamide (**20**). $^1$H NMR (400 MHz, DMSO-$d_6$) δ 8.76 (dd, *J* = 7.8, 4.0 Hz, 1H), 8.37 (d, J = 51.9 Hz 1H), 8.34 (dd, *J* = 12.0, 4.4 Hz, 1H), 7.72 – 7.66 (m, 1H), 7.57 – 7.30 (m, 7H), 7.30 – 7.12 (m, 4H), 7.11 – 7.05 (m, 2 H), 6.38 – 6.30 (m, 2H), 6.23 (dt, *J* = 5.2, 2.2 Hz, 2H), 5.58 (d, *J* = 3.6 Hz, 0.5H), 5.53 (d, *J* = 3.6 Hz, 0.5H), 5.01 (h, *J* = 7.3 Hz, 1H), 1.39 (d, *J* = 7.0 Hz, 1.5H), 1.26 (d, *J* = 7.0 Hz, 1.5H). $^{13}$C NMR (101 MHz, DMSO-$d_6$) δ 168.26, 168.05, 158.28, 158.21,

151.44, 151.37, 148.99, 146.33, 146.27, 145.45, 145.40, 144.29, 143.99, 139.09, 139.08, 137.67, 137.63, 135.76, 135.73, 132.60, 128.24, 128.11, 126.70, 126.65, 126.17, 125.79, 118.73, 118.28, 116.40, 116.35, 111.42, 110.91, 110.89, 61.98, 54.92, 48.40, 48.26, 22.32, 22.16. Yield. 80.3 %. $C_{30}H_{26}N_4O_3$, HRMS calculated for m/z $[M+H]^+$: 491.208315 (calculated), 491.2078 (found).

2-[1-(furan-2-yl)-N-[4-(pyridin-2-yl)phenyl]formamido]-N-[(1S)-1-phenylethyl]-2-(pyridin-3-yl)acetamide (**21**). $^1$H NMR (500 MHz, DMSO-d$_6$) δ 8.77 (dd, *J* = 7.8, 5.8 Hz, 1H), 8.65 – 8.59 (m, 1H), 8.46-8.27 (m, 2H), 7.97 – 7.88 (m, 3H), 7.88 – 7.78 (m, 1H), 7.68 (ddd, *J* = 7.6, 1.7, 0.8 Hz, 1H), 7.57 – 7.00 (m, 10H), 6.36 (d, *J* = 14.4 Hz, 1H), 6.32 (ddd, *J* = 9.0, 3.6, 1.7 Hz, 1H), 5.59 (d, *J* = 3.6, 0.8 Hz, 0.5H), 5.54 (d, *J* = 3.6, 0.8 Hz, 0.5H), 5.09 – 4.94 (m, 1H), 1.39 (d, *J* = 7.0 Hz, 1.5H), 1.27 (d, *J* = 7.0 Hz, 1.5H). $^{13}$C NMR (126 MHz, DMSO-d$_6$) δ 168.22, 168.02, 158.25, 158.19, 154.66, 151.42, 151.36, 149.59, 149.58, 149.01, 148.99, 146.35, 146.30, 145.47, 145.42, 144.32, 144.03, 139.99, 139.95, 138.08, 137.75, 137.72, 137.37, 137.35, 131.68, 130.69, 130.32, 128.28, 128.16, 126.74, 126.69, 126.54, 126.20, 125.83, 123.14, 122.99, 122.85, 120.38, 120.37, 116.44, 116.40, 111.43, 111.41, 62.13, 48.47, 48.33, 22.36, 22.19. Yield. 88.3 %. $C_{31}H_{26}N_4O_3$, HRMS calculated for m/z $[M+H]^+$: 503.208315 (calculated), 503.2078 (found).

2-[N-(4-benzylphenyl)-1-(furan-2-yl)formamido]-N-[(1S)-1-phenylethyl]-2-(pyridin-3-yl)acetamide (**22**). $^1$H NMR (400 MHz, DMSO-d$_6$) δ 8.71 (dd, *J* = 7.8, 4.2 Hz, 1H), 8.44 – 8.23 (m, 2H), 7.66 (ddd, *J* = 5.5, 1.7, 0.7 Hz, 1H), 7.45 – 7.31 (m, 3H), 7.31 – 7.11 (m, 7H), 7.10 – 6.96 (m, 6H), 6.39 – 6.18 (m, 2H), 5.43 – 5.38 (m, 0.5H), 5.36 – 5.30 (m, 0.5H), 5.06 – 4.89 (m, 1H), 3.86 (d, *J* = 6.6 Hz, 2H), 1.37 (d, *J* = 7.0 Hz, 1.5H), 1.25 (d, *J* = 7.0 Hz, 1.5H). $^{13}$C NMR (101 MHz, DMSO-d$_6$) δ δ 168.19, 167.97, 158.21, 158.14, 151.41, 151.37, 148.81, 146.39, 146.33, 145.29, 145.25, 144.29, 143.99, 141.18, 140.98, 137.57, 137.12, 137.09, 131.36, 130.72,

130.37, 129.25, 128.43, 128.40, 128.35, 128.33, 128.21, 128.09, 126.67, 126.61, 126.16, 125.96, 125.94, 125.76, 122.91, 122.61, 116.08, 116.04, 111.19, 62.04, 48.35, 48.22, 40.24, 22.30, 22.11. Yield. 80.1 %. $C_{33}H_{29}N_3O_3$, HRMS calculated for m/z $[M+H]^+$: 516.228717 (calculated), 516.2282 (found).

2-(N-{[1,1'-biphenyl]-4-yl}-1-(furan-2-yl)formamido)-N-[(1S)-1-phenylethyl]-2-(pyridin-3-yl)acetamide (**23**). $^1$H NMR (400 MHz, DMSO-d$_6$) δ 8.75 (dd, *J* = 7.8, 5.0 Hz, 1H), 8.53 – 8.21 (m, 2H), 7.72 – 7.66 (m, 1H), 7.66 – 7.58 (m, 2H), 7.58 – 7.24 (m, 10H), 7.24 – 7.05 (m, 4H), 6.47 – 6.23 (m, 2H), 5.63 – 5.53 (m, 0.5H), 5.52 – 5.43 (m, 0.5H), 5.10 – 4.92 (m, 1H), 1.39 (d, *J* = 7.0 Hz, 1.5H), 1.27 (d, *J* = 7.0 Hz, 1.5H). $^{13}$C NMR (101 MHz, DMSO-d$_6$) δ 168.16, 167.97, 158.24, 158.17, 151.39, 151.33, 148.91, 146.34, 146.28, 145.39, 145.34, 144.29, 143.98, 139.46, 138.55, 138.50, 138.46, 137.64, 131.80, 128.95, 128.93, 128.21, 128.09, 127.84, 126.68, 126.61, 126.53, 126.48, 126.17, 125.78, 116.28, 116.25, 111.35, 62.06, 48.39, 48.25, 22.29, 22.11. Yield. 89.0 %. $C_{32}H_{27}N_3O_3$, HRMS calculated for m/z $[M+H]^+$: 502.213067 (calculated), 502.2132 (found).

2-(N-{[1,1'-biphenyl]-4-yl}-1-(1H-imidazol-4-yl)formamido)-N-[(1S)-1-phenylethyl]-2-(pyridin-3-yl)acetamide (**24**). $^1$H NMR (500 MHz, DMSO-d$_6$) δ 12.76 (s, 1H), 8.74 (d, *J* = 7.9 Hz, 1H), 8.44 (d, *J* = 2.4 Hz, 1H), 8.36 (dd, *J* = 4.8, 1.7 Hz, 1H), 7.73 – 7.32 (br s, 1H), 7.68 – 7.60 (m, 3H), 7.60 – 7.49 (m, 3H), 7.48 – 7.23 (m, 9H), 7.19 (ddd, *J* = 7.8, 4.8, 0.8 Hz, 1H), 6.40 (s, 1H), 5.53 (s, 1H), 5.03 (p, *J* = 7.2 Hz, 1H), 1.26 (d, *J* = 7.0 Hz, 3H). $^{13}$C NMR (126 MHz, DMSO-d$_6$) δ 168.31, 151.32, 148.88, 144.07, 139.61, 138.62, 137.60, 137.03, 132.15, 130.93, 128.97, 128.22, 127.85, 126.66, 126.58, 126.17, 123.03, 62.01, 48.23, 22.16.

Yield. 38.2 %. $C_{31}H_{27}N_5O_2$, HRMS calculated for m/z $[M+H]^+$: 502.224299 (calculated), 502.2238 (found).

2-(N-{[1,1'-biphenyl]-4-yl}-1-(1,2-oxazol-5-yl)formamido)-N-[(1S)-1-phenylethyl]-2-(pyridin-3-yl)acetamide (**25**). $^1H$ NMR (400 MHz, DMSO-$d_6$) δ 8.83 (dd, *J* = 7.7, 5.7 Hz, 1H), 8.56 – 8.26 (m, 3H), 7.65 – 7.45 (m, 5H), 7.44 – 7.06 (m, 11H), 6.35 (d, *J* = 11.2 Hz, 1H), 5.88 – 5.77 (m, 1H), 5.13 – 4.94 (m, 1H), 1.40 (d, *J* = 7.0 Hz, 1.5H), 1.28 (d, *J* = 7.0 Hz, 1.5H). $^{13}C$ NMR (101 MHz, DMSO-$d_6$) δ 167.58, 167.39, 161.72, 161.66, 157.08, 157.01, 151.44, 151.38, 150.78, 149.15, 144.18, 143.87, 139.79, 138.40, 137.67, 137.43, 137.38, 131.43, 130.13, 129.75, 128.96, 128.94, 128.27, 128.14, 127.93, 126.76, 126.70, 126.56, 126.54, 126.16, 125.80, 123.12, 122.81, 107.11, 107.10, 62.38, 48.52, 48.38, 22.26, 22.11. Yield. 82.1 %. $C_{31}H_{26}N_4O_3$, HRMS calculated for m/z $[M+H]^+$: 503.208315 (calculated), 503.2078 (found).

2-(N-{[1,1'-biphenyl]-4-yl}-1-(1,3-oxazol-5-yl)formamido)-N-[(1S)-1-phenylethyl]-2-(pyridin-3-yl)acetamide (**26**). $^1H$ NMR (400 MHz, DMSO-$d_6$) δ 8.80 (dd, *J* = 7.8, 4.7 Hz, 1H), 8.57 – 8.23 (m, 3H), 7.69 – 7.49 (m, 5H), 7.48 – 7.32 (m, 6H), 7.31 – 7.07 (m, 5H), 6.34 (d, *J* = 9.3 Hz, 1H), 5.87 (d, *J* = 16.3 Hz, 1H), 5.02 (h, *J* = 7.6 Hz, 1H), 1.40 (d, *J* = 7.0 Hz, 1.5H), 1.27 (d, *J* = 7.0 Hz, 1.5H). $^{13}C$ NMR (101 MHz, DMSO-$d_6$) δ 167.88, 167.69, 156.94, 156.87, 153.69, 153.65, 151.45, 151.38, 149.10, 144.28, 144.24, 144.21, 143.93, 140.15, 138.45, 137.69, 137.65, 137.51, 137.47, 131.85, 130.41, 130.37, 129.92, 128.99, 128.97, 128.25, 128.13, 127.98, 126.83, 126.73, 126.65, 126.64, 126.16, 125.79, 123.11, 122.81, 62.12, 48.48, 48.34, 22.28, 22.11. Yield. 85.0 %. $C_{31}H_{26}N_4O_3$, HRMS calculated for m/z $[M+H]^+$: 503.208315 (calculated), 503.2078 (found).

N-tert-butyl-2-[N-(4-tert-butylphenyl)-1-(furan-2-yl)formamido]-2-(pyrazin-2-yl)acetamide (**27**). $^{1}$H NMR (500 MHz, DMSO-$d_6$) δ 8.53 (dd, *J* = 2.5, 1.5 Hz, 1H), 8.47 (d, *J* = 1.5 Hz, 1H), 8.43 (d, *J* = 2.6 Hz, 1H), 8.00 (s, 1H), 7.68 (dd, *J* = 1.7, 0.7 Hz, 1H), 7.35 – 7.16 (m, 4H), 6.33 (dd, *J* = 3.6, 1.7 Hz, 1H), 6.28 (s, 1H), 5.39 (d, *J* = 3.6 Hz, 1H), 1.22 (s, 9H), 1.14 (s, 9H). $^{13}$C NMR (126 MHz, DMSO-$d_6$) δ 166.07, 158.30, 152.04, 151.03, 146.24, 145.34, 145.28, 143.63, 143.03, 136.93, 130.26, 125.29, 116.05, 111.29, 65.10, 50.51, 34.31, 31.02, 28.17.
Yield. 86.1 %. $C_{25}H_{30}N_4O_3$, HRMS calculated for m/z $[M+H]^+$: 435.239615 (calculated), 435.2391 (found).

N-tert-butyl-2-[N-(4-tert-butylphenyl)-1-(furan-2-yl)formamido]-2-(1H-imidazol-4-yl)acetamide (**28**). $^{1}$H NMR (400 MHz, DMSO-$d_6$) δ 8.91 (d, *J* = 1.3 Hz, 1H), 8.02 (s, 1H), 7.73 – 7.66 (m, 1H), 7.36 – 7.29 (m, 2H), 7.23 (dd, *J* = 1.3, 0.7 Hz, 3H), 6.34 (dd, *J* = 3.6, 1.7 Hz, 1H), 6.27 (s, 1H), 5.34 (d, *J* = 3.6 Hz, 1H), 1.24 (s, 9H), 1.23 (s, 9H). $^{13}$C NMR (101 MHz, DMSO-$d_6$) δ 165.47, 158.23, 151.48, 146.02, 145.56, 136.40, 134.59, 129.92, 128.33, 125.57, 119.68, 116.29, 111.41, 56.24, 50.78, 34.40, 31.04, 28.24. Yield. 60.4 %. $C_{24}H_{30}N_4O_3$, HRMS calculated for m/z $[M+H]^+$: 423.239615 (calculated), 423.2391 (found).

N-tert-butyl-2-[N-(4-tert-butylphenyl)-1-(furan-2-yl)formamido]-2-(pyrimidin-5-yl)acetamide (**29**). $^{1}$H NMR (400 MHz, DMSO-$d_6$) δ 8.93 (s, 1H), 8.46 (s, 2H), 7.95 (s, 1H), 7.68 (dd, *J* = 1.7, 0.8 Hz, 1H), 7.33-7.12 (br s, 2H)7.27 (d, *J* = 8.0 Hz, 2H), 6.32 (dd, *J* = 3.6, 1.7 Hz, 1H), 6.16 (s, 1H), 5.32 (d, *J* = 3.6 Hz, 1H), 1.22 (s, 9H), 1.21 (s, 9H). $^{13}$C NMR (101 MHz, DMSO-$d_6$) δ 167.13, 158.13, 157.87, 157.29, 151.36, 146.22, 145.36, 136.41, 130.79, 129.64, 125.58, 116.01,

111.31, 60.53, 50.59, 34.33, 30.98, 28.23. Yield. 78.4 %. $C_{25}H_{30}N_4O_3$, HRMS calculated for m/z $[M+H]^+$: 435.239615 (calculated), 435.2391 (found).

2-(N-{[1,1'-biphenyl]-4-yl}-1-(furan-2-yl)formamido)-N-cyclopropyl-2-(pyridin-3-yl)acetamide (**30**). $^1$H NMR (500 MHz, DMSO-$d_6$) δ 8.46 – 8.26 (m, 3H), 7.69 (dd, *J* = 1.7, 0.8 Hz, 1H), 7.66 – 7.62 (m, 2H), 7.62 – 7.50 (m, 2H), 7.49-7.14 (br s, 1H), 7.48 – 7.31 (m, 5H), 7.17 (dd, *J* = 7.9, 4.8 Hz, 1H), 6.34 (dd, *J* = 3.6, 1.7 Hz, 1H), 6.18 (s, 1H), 5.55 (dd, *J* = 3.6, 0.8 Hz, 1H), 2.74 – 2.65 (m, 1H), 0.73 – 0.55 (m, 2H), 0.46 – 0.26 (m, 2H). $^{13}$C NMR (126 MHz, DMSO-$d_6$) δ 169.94, 158.22, 151.19, 148.91, 146.31, 145.41, 139.50, 138.56, 138.52, 137.54, 131.72, 130.62, 128.96, 127.86, 126.55, 126.53, 123.04, 116.32, 111.37, 62.06, 22.51, 5.59, 5.57.
Yield. 89.8 %. $C_{27}H_{23}N_3O_3$, HRMS calculated for m/z $[M+H]^+$: 438.181767 (calculated), 438.1812 (found).

2-(N-{[1,1'-biphenyl]-4-yl}-1-(furan-2-yl)formamido)-2-(pyridin-3-yl)-N-(2,4,4-trimethylpentan-2-yl)acetamide (**31**). $^1$H NMR (400 MHz, DMSO-$d_6$) δ 8.48 – 8.22 (m, 2H), 7.75 – 7.66 (m, 2H), 7.66 – 7.60 (m, 2H), 7.60 – 7.49 (m, 2H), 7.51-7.09 (br s, 1H), 7.49 – 7.30 (m, 5H), 7.15 (ddd, *J* = 7.9, 4.8, 0.8 Hz, 1H), 6.32 (dd, *J* = 3.6, 1.7 Hz, 1H), 6.25 (s, 1H), 5.49 (dd, *J* = 3.6, 0.8 Hz, 1H), 1.68 (dd, *J* = 75.4, 14.5 Hz, 2H), 1.29 (d, *J* = 16.4 Hz, 6H), 0.87 (s, 9H). $^{13}$C NMR (101 MHz, DMSO-$d_6$) δ 167.73, 158.09, 151.31, 148.69, 146.39, 145.25, 139.43, 138.69, 138.61, 137.61, 131.90, 130.93, 128.95, 127.83, 126.53, 126.46, 122.81, 116.11, 111.33, 62.43, 50.71, 31.19, 31.11, 29.03, 28.40. Yield. 62.1 %. $C_{32}H_{35}N_3O_3$, HRMS calculated for m/z $[M+H]^+$: 510.275667 (calculated), 510.2751 (found).

2-(N-{[1,1'-biphenyl]-4-yl}-1-(furan-2-yl)formamido)-N-cyclopentyl-2-(pyridin-3-yl)acetamide (**32**). $^1$H NMR (400 MHz, DMSO-$d_6$) δ 8.35 (d, *J* = 20.6 Hz, 2H), 8.24 (d, *J* = 7.0 Hz, 1H), 7.68

(dd, *J* = 1.7, 0.7 Hz, 1H), 7.66 – 7.60 (m, 2H), 7.55 (d, *J* = 8.1 Hz, 2H), 7.51-7.12 (br s, 1H), 7.50 – 7.29 (m, 5H), 7.16 (dd, *J* = 7.9, 4.7 Hz, 1H), 6.33 (dd, *J* = 3.6, 1.7 Hz, 1H), 6.24 (s, 1H), 5.53 (d, *J* = 3.6 Hz, 1H), 4.06 (h, *J* = 6.6 Hz, 1H), 1.90 – 1.70 (m, 2H), 1.68 – 1.37 (m, 5H), 1.33 – 1.19 (m, 1H). $^{13}$C NMR (101 MHz, DMSO-d$_6$) δ 168.21, 158.22, 151.22, 148.83, 146.36, 145.36, 139.46, 138.58, 137.52, 131.78, 130.92, 128.97, 127.85, 126.55, 126.49, 123.02, 116.26, 111.37, 62.07, 50.75, 32.13, 31.87, 23.49, 23.44. Yield. 78.8 %. $C_{29}H_{27}N_3O_3$, HRMS calculated for m/z $[M+H]^+$: 466.213067 (calculated), 466.2125 (found).

2-(N-{[1,1'-biphenyl]-4-yl}-1-(1H-imidazol-4-yl)formamido)-N-cyclopentyl-2-(pyridin-3-yl)acetamide (**33**). $^1$H NMR (500 MHz, DMSO-d$_6$) δ 8.38 (d, *J* = 2.3 Hz, 1H), 8.33 (dd, *J* = 4.8, 1.7 Hz, 1H), 8.24 (d, *J* = 7.1 Hz, 1H), 7.82-7.32 (br s, 1H), 7.73 – 7.55 (m, 5H), 7.53 – 7.29 (m, 5H), 7.23 – 7.12 (m, 1H), 6.24 (s, 1H), 5.54 (s, 1H), 4.12 – 4.00 (m, 1H) 1.90 – 1.70 (m, 2H), 1.67 – 1.40 (m, 5H), 1.32 – 1.19 (m, 1H). $^{13}$C NMR (126 MHz, DMSO-d$_6$) δ 168.46, 151.23, 148.83, 139.65, 138.74, 138.67, 137.56, 132.17, 131.10, 129.04, 127.91, 126.63, 123.05, 62.09, 50.73, 32.14, 31.99, 23.54, 23.48. Yield. 44.8 %. $C_{28}H_{27}N_5O_2$, HRMS calculated for m/z $[M+H]^+$: 466.224299 (calculated), 466.2238 (found).

2-(N-{[1,1'-biphenyl]-4-yl}-1-(furan-2-yl)formamido)-N-cyclohexyl-2-(pyridin-3-yl)acetamide (**34**). $^1$H NMR (500 MHz, $CDCl_3$/ MeOD-d$_4$) δ 8.43 (s, 1H), 8.38 – 8.31 (m, 1H), 7.55 – 7.48 (m, 3H), 7.48 – 7.27 (m, 6H), 7.26 – 6.95 (br s, 1H), 7.16 – 7.05 (m, 2H), 6.22 (s, 1H), 6.16 – 6.08 (m, 1H), 5.55 (d, *J* = 3.6 Hz, 1H), 3.79 (s, 1H), 3.77 – 3.69 (m, 1H), 2.04 – 1.46 (m, 5H), 1.43 – 0.9 (m, 5H). $^{13}$C NMR (126 MHz, $CDCl_3$/ MeOD-d$_4$) δ 168.11, 168.03, 159.83, 150.90, 148.99, 145.91, 145.15, 141.54, 139.33, 138.51, 137.96, 131.38, 130.82, 128.81, 127.86, 127.41, 126.89, 123.32, 117.57, 111.32, 62.70, 62.66, 32.51, 32.47, 25.35, 24.78, 24.71.

Yield. 83.8 %. $C_{30}H_{29}N_3O_3$, HRMS calculated for m/z $[M+H]^+$: 480.228716 (calculated), 480.2282 (found).

2-(N-{[1,1'-biphenyl]-4-yl}-1-(1H-imidazol-4-yl)formamido)-N-cyclohexyl-2-(pyridin-3-yl)acetamide (**35**). $^1$H NMR (500 MHz, DMSO-d$_6$) δ 8.40 (d, *J* = 2.3 Hz, 1H), 8.33 (dd, *J* = 4.8, 1.7 Hz, 1H), 8.17 (d, *J* = 7.7 Hz, 1H), 7.89 (s, 1H), 7.69 – 7.63 (m, 2H), 7.59 (d, *J* = 7.2 Hz, 2H), 7.54-6.99 (br s, 1 H), 7.51 – 7.32 (m, 5H), 7.17 (ddd, *J* = 7.9, 4.8, 0.8 Hz, 1H), 6.26 (s, 1H), 5.56 (s, 1H), 3.67-3.57 (m, 1H), 1.86 – 1.48 (m, 5H), 1.35 – 1.16 (m, 3H), 1.15 – 0.96 (m, 2H). $^{13}$C NMR (126 MHz, DMSO-d$_6$) δ 167.86, 159.86, 151.22, 148.81, 139.81, 138.59, 138.44, 137.65, 136.86, 132.09, 130.99, 129.02, 127.94, 126.71, 126.63, 123.07, 62.18, 48.03, 32.18, 25.18, 24.56, 24.41. Yield. 42.8 %. $C_{29}H_{29}N_5O_2$, HRMS calculated for m/z $[M+H]^+$: 480.239950 (calculated), 480.2394 (found).

2-[N-(4-cyclohexylphenyl)-1-(1,2-oxazol-5-yl)formamido]-N-[(1S)-1-phenylethyl]-2-(pyridin-3-yl)acetamide (**36**). $^1$H NMR (400 MHz, DMSO-d$_6$) δ 8.69 (dd, *J* = 7.8, 3.1 Hz, 1H), 8.32 (ddd, *J* = 10.3, 4.8, 1.6 Hz, 1H), 8.31 (dd, J = 46.3, 2.2 Hz, 1H), 7.74 – 7.60 (m, 1H), 7.46 – 7.29 (m, 3H), 7.28 – 6.96 (m, 7H), 6.35 – 6.19 (m, 2H), 5.26 – 5.22 (m, 0.5H), 5.21 – 5.15 (m, 0.5H), 5.08 – 4.87 (m, 1H), 2.48 – 2.34 (m, 1H), 1.87 – 1.59 (m, 5H), 1.47 – 1.07 (m, 5H), 1.31 (dd, J = 47.0 7.0 Hz, 3 H) $^{13}$C NMR (101 MHz, DMSO-d$_6$) δ 168.20, 167.97, 158.25, 158.16, 151.35, 151.30, 148.80, 147.89, 146.34, 146.28, 145.30, 145.26, 144.32, 144.02, 137.53, 136.69, 136.66, 131.14, 130.77, 130.44, 128.21, 128.08, 126.74, 126.67, 126.60, 126.17, 125.76, 122.86, 122.56, 115.89, 115.85, 111.26, 62.03, 54.92, 48.35, 48.22, 43.11, 33.86, 33.81, 33.78, 26.19, 25.48, 22.29,

22.11. Yield. 76.3 %. $C_{31}H_{32}N_4O_3$, HRMS calculated for m/z $[M+H]^+$: 509.255266 (calculated), 509.2547 (found).

2-(N-{[1,1'-biphenyl]-4-yl}-1-(1H-imidazol-4-yl)formamido)-N-[(1S)-1-phenylethyl]-2-(pyrazin-2-yl)acetamide (**37**). $^1$H NMR (500 MHz, DMSO-$d_6$) δ 8.89 – 8.37 (m, 5H), 7.70 – 7.61 (m, 3H), 7.57 (dd, *J* = 16.7, 8.1 Hz, 2H), 7.52 – 7.41 (m, 3H), 7.40 – 7.29 (m, 4H), 7.29 – 7.16 (m, 3H), 6.61 – 6.47 (m, 1H), 5.07 – 4.93 (m, 1H), 1.33 (d, *J* = 7.0 Hz, 1.5H), 1.30 (d, *J* = 7.0 Hz, 1.5H). $^{13}$C NMR (126 MHz, DMSO-$d_6$) δ 166.72, 166.57, 151.45, 151.33, 146.17, 146.00, 144.20, 144.15, 143.83, 143.78, 143.44, 138.74, 131.67, 131.59, 129.04, 128.25, 128.18, 127.91, 126.69, 126.65, 126.16, 126.02, 64.53, 48.44, 48.39, 22.24, 22.22. Yield. 42.3 %. $C_{30}H_{26}N_6O_2$, HRMS calculated for m/z $[M+H]^+$: 503.219549 (calculated), 503.2190 (found).

2-(N-{[1,1'-biphenyl]-4-yl}-1-(furan-2-yl)formamido)-N-[(1S)-1-phenylethyl]-2-(pyrazin-2-yl)acetamide (**38**). $^1$H NMR (500 MHz, DMSO-$d_6$) δ 8.79 (dd, *J* = 24.5, 7.8 Hz, 1H), 8.68 – 8.34 (m, 3H), 7.72 – 7.67 (m, 1H), 7.66 – 7.61 (m, 2H), 7.60 – 7.51 (m, 2H), 7.49-7.26 (br s, 1H), 7.48 – 7.41 (m, 2H), 7.39 – 7.30 (m, 4H), 7.29 – 7.16 (m, 3H), 6.56 – 6.46 (m, 1H), 6.40 – 6.31 (m, 1H), 5.66 – 5.64 (m, 0.5H), 5.62 – 5.59 (m, 0.5H), 5.10 – 4.90 (m, 1H), 1.39 – 1.24 (m, 3H). $^{13}$C NMR (126 MHz, DMSO-$d_6$) δ 166.52, 166.33, 158.39, 158.32, 151.22, 151.10, 146.21, 146.17, 146.06, 145.53, 145.49, 144.16, 144.05, 143.83, 143.77, 143.47, 139.63, 139.57, 138.88, 138.83, 138.67, 138.66, 131.23, 131.16, 129.02, 128.25, 128.18, 127.90, 126.70, 126.61, 126.16, 126.04, 116.62, 116.59, 111.47, 111.44, 64.50, 64.46, 48.51, 48.46, 22.22. Yield. 83.2 %. $C_{31}H_{26}N_4O_3$, HRMS calculated for m/z $[M+H]^+$: 503.208315 (calculated), 503.2078 (found).

2-(N-{[1,1'-biphenyl]-4-yl}-1-(furan-2-yl)formamido)-N-(2,2-diphenylethyl)-2-(pyrazin-2-yl)acetamide (**39**). $^{1}$H NMR (500 MHz, DMSO-d$_6$) δ 8.39 – 8.19 (m, 3H), 7.69 (dd, *J* = 1.7, 0.7 Hz, 1H), 7.67 – 7.60 (m, 2H), 7.58 – 7.50 (m, 2H), 7.46 – 7.42 (m, 2H), 7.38 – 7.33 (m, 1H), 7.32 – 7.10 (m, 12H), 7.05 (dt, *J* = 7.9, 2.0 Hz, 1H), 6.96 (dd, *J* = 8.0, 4.8 Hz, 1H), 6.33 (dd, *J* = 3.6, 1.7 Hz, 1H), 6.21 (s, 1H), 5.52 (dd, *J* = 3.6, 0.8 Hz, 1H), 4.23 (t, *J* = 7.9 Hz, 1H), 4.06 – 3.90 (m, 1H), 3.73 – 3.57 (m, 1H). $^{13}$C NMR (126 MHz, DMSO-d$_6$) δ 168.79, 158.24, 151.45, 148.59, 146.33, 145.39, 142.79, 142.65, 139.45, 138.56, 138.55, 137.45, 131.63, 130.52, 129.00, 128.40, 128.38, 127.92, 127.88, 127.85, 126.56, 126.52, 126.34, 126.33, 122.78, 116.34, 111.38, 62.13, 50.06, 43.60. Yield. 65.5 %. $C_{38}H_{31}N_3O_3$, HRMS calculated for m/z [M+H]$^+$: 578.244367 (calculated), 578.2438 (found).

2-(N-{[1,1'-biphenyl]-4-yl}-1-(furan-2-yl)formamido)-N-(3-phenylpropyl)-2-(pyridin-3-yl)acetamide (**40**). $^{1}$H NMR (500 MHz, DMSO-*d$_6$*) δ 8.55 (d, *J* = 2.2 Hz, 1H), 8.49 (dd, *J* = 5.2, 1.6 Hz, 1H), 8.32 (t, *J* = 5.5 Hz, 1H), 7.89 – 7.79 (m, 1H), 7.64 (d, *J* = 1.5 Hz, 1H), 7.63 – 7.51 (m, 4H), 7.49 – 7.42 (m, 1H), 7.42 – 7.21 (m, 5H), 7.21 – 7.13 (m, 2H), 7.12 – 7.06 (m, 3H), 6.30 (dd, *J* = 3.6, 1.7 Hz, 1H), 6.19 (s, 1H), 5.58 (d, *J* = 3.5 Hz, 1H), 3.08 (q, *J* = 6.6 Hz, 2H), 2.48 – 2.39 (m, 2H), 1.69 – 1.55 (m, 2H). $^{13}$C NMR (101 MHz, DMSO-*d$_6$*) δ 167.76, 158.33, 146.18, 145.57, 141.68, 139.76, 138.89, 138.55, 132.80, 131.28, 128.98, 128.28, 128.26, 127.92, 126.86, 126.60, 125.71, 124.53, 116.67, 111.46, 62.67, 38.63, 32.44, 30.78. Yield: 76.5 %. $C_{33}H_{29}N_3O_3$, HRMS calculated for m/z [M+H]$^+$: 516.228717 (calculated), 516.2282 (found).

(2R)-2-(N-{[1,1'-biphenyl]-4-yl}-1-(furan-2-yl)formamido)-N-[(1S)-1-phenylethyl]-2-(pyridin-3-yl)acetamide (**23R**). $^{1}$H NMR (500 MHz, DMSO-*d$_6$*) δ 8.79 (d, *J* = 7.9 Hz, 1H), 8.64 (d, *J* = 2.0 Hz, 1H), 8.56 (d, *J* = 5.2 Hz, 1H), 7.87 (d, *J* = 8.0 Hz, 1H), 7.69 (s, 1H), 7.64 (d, *J* = 7.7 Hz,

2H), 7.57 (d, *J* = 8.2 Hz, 2H), 7.54 – 7.48 (m, 1H), 7.48 – 7.12 (m, 10H), 6.41 (s, 1H), 6.38 – 6.27 (m, 1H), 5.59 (d, *J* = 3.6 Hz, 1H), 5.02 (t, *J* = 7.2 Hz, 1H), 1.30 (d, *J* = 7.0 Hz, 3H). $^{13}$C NMR (126 MHz, DMSO) δ 167.30, 158.27, 147.98, 146.17, 145.85, 145.52, 143.81, 141.61, 139.71, 138.56, 138.53, 132.66, 131.52, 128.97, 128.24, 127.92, 126.75, 126.59, 126.21, 124.54, 116.56, 111.43, 62.05, 48.39, 21.98. $C_{32}H_{27}N_3O_3$, HRMS calculated for m/z $[M+H]^+$: 502.213067 (calculated), 502.2125 (found).

(2S)-2-(N-{[1,1'-biphenyl]-4-yl}-1-(furan-2-yl)formamido)-N-[(1S)-1-phenylethyl]-2-(pyridin-3-yl)acetamide (**23S**). $^{1}$H NMR (500 MHz, DMSO-*d*$_6$) δ 8.82 (d, *J* = 7.7 Hz, 1H), 8.53 (s, 2H), 7.79 – 7.51 (m, 6H), 7.51 – 7.05 (m, 11H), 6.39 (s, 1H), 6.36 (dd, *J* = 3.5, 1.7 Hz, 1H), 5.64 (d, *J* = 3.5 Hz, 1H), 4.98 (p, *J* = 7.1 Hz, 1H), 1.37 (d, *J* = 7.0 Hz, 3H). $^{13}$C NMR (126 MHz, DMSO) δ 167.13, 158.33, 148.00, 146.21, 145.86, 145.56, 144.14, 141.52, 139.74, 138.55, 138.51, 132.36, 131.56, 128.97, 128.15, 127.91, 126.76, 126.68, 126.59, 125.83, 124.28, 116.59, 111.45, 62.02, 48.55, 22.18. $C_{32}H_{27}N_3O_3$, HRMS calculated for m/z $[M+H]^+$: 502.213067 (calculated), 502.2125 (found).
